# Hierarchical diversification of neurons regulating motivated behaviors

**DOI:** 10.1101/2025.06.03.657692

**Authors:** Najia A. Elkahlah, Yunzhi Lin, Yijie Pan, Joseph A. Carter, Troy R. Shirangi, E. Josephine Clowney

## Abstract

Brain regions that regulate motivated behaviors, including the vertebrate hypothalamus and arthropod cerebrum, house bespoke neural circuits dedicated to perceptual and internal regulation of many behavioral states^1,2^. These circuits are built to purpose from complex sets of cell types whose patterning has been challenging to elucidate. Here, we developed methods in *Drosophila melanogaster* to embed well-studied neurons that regulate mating in the transcriptional contexts of the neuronal lineages that generate them^3–5^. By comparing transcription within and between lineages, we identified a large set of transcription factors expressed in complex combinations that delineate cerebral hemilineages – classes of postmitotic neurons born from the same stem cell and sharing Notch status^6,7^. Hemilineages comprise the major anatomic classes in the cerebrum^8–10^ and these transcription factors are required to generate their gross features. We show that subtypes of the same hemilineage can provide a common computational module to circuits regulating different drives, and identify an orthogonal set of transcription factors that stratify hemilineage subtypes of differing birth order. Our findings suggest that distinct sets of transcription factors operate in a hierarchical system to build, diversify, and sexually differentiate lineally-related neurons that compose motivated behaviors circuits. By linking developmental patterning to separable transcriptional axes that produce gross versus fine aspects of information flow, we provide a logical framework for cerebral control of diverse drives.

## Introduction

Motivated behaviors, like mating, feeding, or aggression, are regulated by neurons built to purpose for singular information-processing functions^1,2^. These wire into unique feed-forward circuits in vertebrate subcortical nuclei and the arthropod cerebrum. The genome contains the information to build meanings of sensory stimuli and relationships among drives into the structures of motivated behavior circuits, yet their immense diversity has made their patterning too difficult to address. To explicate anatomic and functional principles structuring these circuits and states, we need to describe the logical principles and molecular mechanisms through which their constituent neurons are patterned during development.

Four logical axes diversify neuronal cell types; they are used distinctly across brain regions within organisms and in different kinds of neurogenesis programs across clades^11–13^: Neural stem cells are (1) spatially differentiated, and (2) progress through temporal expression windows as they divide asymmetrically. Differentiating daughter cells can be further separated by (3) a Notch switch, and (4) sex further differentiates a subset of cell types.

In the arthropod cerebrum, ∼200 anatomic units called hemilineages form an information highway system traversing brain regions, as their constituent neurons share neurite outgrowth tracts, gross morphology, and, often, neurotransmitter systems^8–10,14–17^. In adult *Drosophila melanogaster*, cerebral hemilineages, like those of the ventral nerve cord (VNC), are composed of the NotchON or NotchOFF cousins from the same neuroblast^7–10,18,19^. Within hemilineage groundplans, individual cells have identifiable differences in fine axonal and dendritic anatomy and connectivity, linked to their birth order^15,18,20,21^. While lineages of nerve cords, visual systems, or cortex repeat across segments or columns, the unique functions of the cerebrum are produced by mostly singular lineages. Thus cerebral “cell types” defined at the level of anatomy, connectivity, or circuit function are often specific birth order cohorts from within one hemilineage. This singularity is what makes the complex circuit computations of the cerebrum possible; at the same time, the rarity of these neuroblasts and their daughter cells has impeded efforts to describe the transcriptional mechanisms that link neurogenic patterning to the formation of circuit architecture.

Here, we break this impasse by starting from the unusually well-understood circuit that regulates male mating in *Drosophila melanogaster*. This circuit is built from >60 populations of neurons that arise as subtypes of defined lineages and that are sexually differentiated by the Fruitless and/or Doublesex transcription factors^3,4,10,18,22,23^. We transcriptionally characterized neuronal subtypes in the mating circuit in the context of their lineages and identified groups of transcription factors that play separable logical roles in diversifying hemilineage ground plans, versus birth-order-associated subtypes within hemilineages. Focusing on a hemilineage that generates sexually differentiated neurons that gate mating, we found that mutation of hemilineage TFs disrupted general morphological features; that manipulation of birth order TFs shifted subtype fates; and that cellular sex differentiated transcription within subtypes.

Our results demonstrate that cerebral neurons with singular circuit functions are specified through a nested, hierarchical code: hemilineage identity sets the ground plan, while birth order carves this plan into distinct subtypes and sexual differentiation overlays additional specialization onto selected cohorts. These hierarchically arranged bits of transcriptional information durably barcode neurons, allowing the brain to build uniquely purposed circuit elements from the same basic developmental logic.

## Results

### Transcriptional analysis of developing cerebral lineages

Most of the adult *Drosophila melanogaster* cerebrum derives from ∼100 spatially patterned, bilaterally symmetric Type I neuroblasts, which each divide 70-100 times from embryonic to pupal stages as they progress through expression of temporal transcription factors (TFs) and RNA binding proteins^24^. Neuroblast asymmetric divisions produce a ganglion mother cell, and these divide once to produce two neurons, who assume NotchON versus NotchOFF fates^6,7,25^. The 70-100 neurons of the same neuroblast that share Notch status form a hemilineage. To interrogate hemilineage patterning in the cerebrum, we used an intersectional genetic approach to lineage trace the progeny of identified cerebral neuroblasts that produce well-studied, *fruitless*-expressing neurons in the male mating circuit (Fig. 1a). In this system, expression of GAL4 within the neuroblast initiates a recombination cascade, resulting in permanent expression of LexA within its progeny (Fig. 1b)^5^. We drove lineage tracing using the R19C05*^Pdfr^* enhancer, which has been empirically shown to be expressed in the CREa1 and SMPad1 neuroblasts, among additional lineages (Figure 1b, d)^22^. The NotchOFF (B) hemilineage from CREa1 includes *fruitless* mAL neurons, which inhibit courtship-promoting P1 neurons in the presence of males or heterospecific females^18,26–28^. The NotchON (A) hemilineage from SMPad1 includes *fruitless* aSP-2 neurons, which are activated in the presence of females and enhance courtship persistence^3,4,10,29^. We registered twenty labeled brains to the FAFB connectome and annotated 13 lineages labeled in at least half of hemispheres, and 13 additional clones labeled less frequently (Fig. 1d; Extended Data Fig. 2; Methods). Using this method, all neurons born after the completion of the recombination cascade are labeled, thus embryonically-born (primary) neurons may not be included.

**Figure 1.**
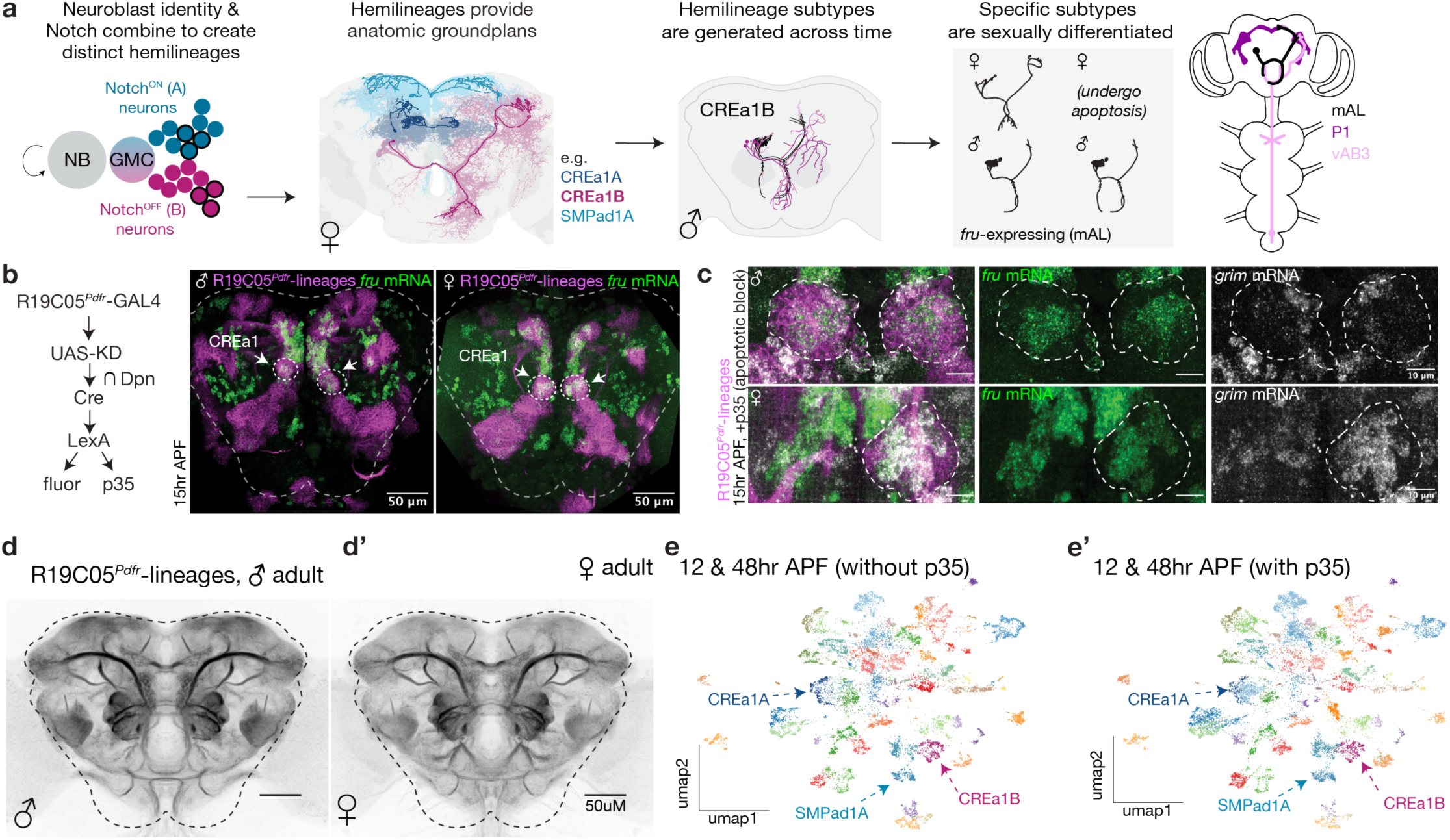
Transcriptional analysis of developing cerebral lineages. **a**, Type I neuroblasts generate hemilineages, which form anatomic highways structuring arthropod cerebral circuits. The *D. melanogaster* CREa1B hemilineage produces *fruitless* mAL neurons (black) and other morphologies across time. Right: mAL neurons receive gustatory pheromone input from vAB3 and inhibit P1 neurons to gate courtship^15,18,26,27^. **b**, Intersection of R19C05*^Pdfr^* enhancer with *deadpan* allows lineage tracing of CREa1 and other NBs (magenta)^5,22^, including their *fruitless* progeny (RNA FISH, green). Dashed lines outline central brain. Additional information provided in Extended Data Figure 1 and Methods. **c**, Clonal induction of p35 in the CREa1 lineage prevents the completion of apoptosis. CREa1B soma marked by R19C05*^Pdfr^*-lineage labeling, combined with RNA FISH for *fru* and *grim*. *grim* transcript marks neurons fated to apoptose. In the female sample shown, CREa1B is labeled (and p35 expressed) only in the right hemisphere. In male, *fru* is translated into FruM and suppresses *grim*. **d**, Registered, average signal of 10 male or female R19C05*^Pdfr^* lineage-traced brains, highlighting hemilineage tract morphologies. Census of labeled lineages provided in Extended Data Figure 2. **e**, **e’**, We performed scRNAseq transcriptional analysis of developing R19C05*^Pdfr^* lineage neurons (12 and 48h APF), with and without the p35 apoptosis block. We clustered neurons using transcription factor transcripts. QC and analysis details provided in Extended Data Figure 1, and CREa1A, CREa1B and SMPad1A hemilineage markers in Extended Data Figure 2. Here and throughout, allele and full genotype tables are provided in the Methods.

Some neuroblasts produce two living hemilineages with distinct ground plan and neurotransmitter usage, while others produce one living hemilineage and one that undergoes programmed cell death in part or in full^6,7,18,30^. Sexually differentiated cell death is also common^26^. We therefore genetically blocked the completion of apoptosis by expressing the caspase inhibitor p35 using our lineage labeling strategy (Fig. 1b,c)^22^. Resurrected cells were easily identified in histology by their enduring high-level transcription of the pro-apoptotic gene *grim* (Fig. 1c). Using prior clonal analyses, connectomic annotations, and comparative histology in brains with and without p35, we confirmed and updated which pairs of living hemilineages are sisters, and which are lone hemilineages, with apoptosing sisters (Methods, Supplementary Table 1).

Having established the ground truth of the lineages included in our genotype, we FAC-sorted labeled neurons from both sexes, with and without p35, and performed scRNA-seq at 12hr <u>A</u>fter <u>P</u>uparium <u>F</u>ormation (around the end of neurogenesis) and 48hrAPF (when neurites have reached target neuropils and are beginning to contact potential partners) (Extended Data Fig. 1). We retained central brain neurons and sexed them post-hoc using expression of the male-specific lncRNAs *rox1* and *rox2* (Methods, Extended Data Fig. 1). To study the diversification of neuronal fates, we performed semi-supervised clustering using the 628 annotated *D. mel* transcription factors. After integrating time points, we tested a variety of methods and resolutions to probe how the four logical axes of diversity related to patterns of transcription factor expression in the dataset and selected an intermediate resolution based on standard bioinformatic tools, producing 56 clusters after further manual curation (Methods, Fig. 1e, Extended Data Fig. 1). At this resolution, neurons did not separate by sex, suggesting that lineage, Notch status, and birth order dominate neuronal patterning^31^. “Resurrected” cells (i.e. with high levels of pro-apoptotic genes) derived only from libraries in which we blocked the completion of apoptosis (Extended Data Fig. 2). The majority of resurrected cells co-clustered with natural cells, but a few clusters were dominated by resurrected cells, indicating potentially novel cell types. We also observed novel neuronal tracts in these animals (Extended Data Fig. 2).

We next used known markers to match neuron types within the R19C05*^Pdfr^* lineages in transcriptional space: mAL neurons from CRE1aB, mushroom body dopaminergic neurons from CREa1A (PAM-DANs), and aSP-2 neurons from SMPad1A (Extended Data Fig. 2). While we didn’t use neurotransmitter transcripts in clustering, we found that most clusters in our dataset expressed unified neurotransmitter systems, as do hemilineages^14^. Moreover, *fruitless* neuron types in our dataset that are subsets of known hemilineages formed subsets of transcriptional clusters that also included non-*fruitless* neurons (Extended Data Fig. 2). We therefore adopted the working hypothesis that our transcriptional clusters correspond to hemilineages, rather than single cell types. If this is the case, then hemilineage [lineage x Notch status] produces the primary variation in transcription factor expression across developing cerebral neurons. Intriguingly, this is consistent with the idea that hemilineages provide the major anatomic building blocks of the arthropod brain and ventral nerve cord^8,17,19^.

### Distinct transcription factor combinations define two sister hemilineages

We used the CREa1 lineage, which produces GABAergic mAL neurons within the NotchOFF B hemilineage and PAM-DANs within the NotchON A hemilineage^18^, as a model to test the hypothesis that our transcriptional clusters represent anatomic hemilineages, with these *fru* subtypes embedded in them. Seven TFs were enriched and pervasive in putative CREa1B cells and endured across developmental time: *Fer1* (bHLH family, homologue of vertebrate PTF1A)*, TfAP-2* (AP-2 family, AP-2 homologue)*, fd59A* (forkhead, FoxD3 homologue)*, dsf* (nuclear receptor, NR2E1/TLX homologue)*, tup* (LIM homeobox, ISL1 homologue) and *ara* and *caup* (both TALE homeobox, homologous to IRX4) (Fig. 2a, b, Extended Data Fig. 3). These TFs were all excluded from the putative CREa1A cluster, which was instead marked by *scro*, *dmrt99B*, *erm*, *D, Lim1*, *hth, SoxN, Fer2 and CG9932* (Extended Data Fig. 4). Though putative CREa1A or CREa1B TFs were not shared between them, each was expressed in other clusters in our transcriptional dataset (Extended Data Fig. 3, 4). The presence of *dsf* in CREa1B and *Fer2* in CREa1A match prior studies^32,33^. Notably, *dsf* mutant males court promiscuously, consistent with a requirement for Dsf in mAL neuron function^34^.

**Figure 2.**
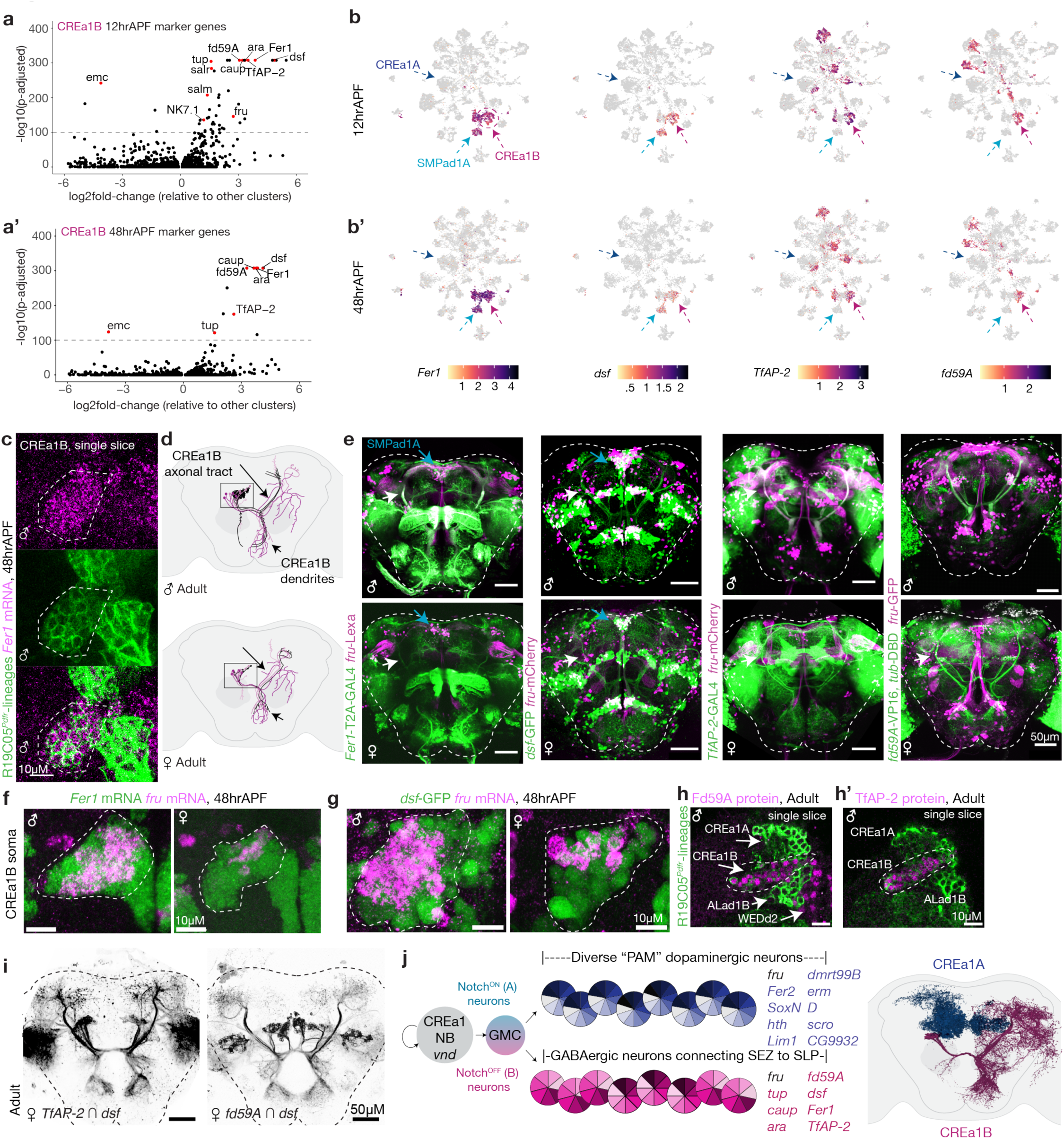
A unique transcription factor combination defines a model hemilineage. **a**, Volcano plots showing differential gene expression between putative CREa1B and other clusters in the dataset identifies hemilineage candidate factors at 12 and 48hr APF. Labeled genes: TFs with -log10(p-adjusted) > 100 based on Wilcoxon Rank Sum test. CREa1A factors shown in Extended Data Figure 3. **b**, **b’**, UMAP plots of putative hemilineage-defining TFs in scRNAseq data from the integrated R19C05*^Pdfr^*-labeled lineages split by time. **c**, *Fer1* RNA fills the CREa1B lineage clone (dashed outline). **d**, CREa1B neuronal morphologies in male and female brains; black are *fru*-expressing mAL neurons. Morphologies informed by^15,18,26^. **e**, Confocal or two-photon Images of cerebral neurons expressing *Fer1*, *dsf*, *TfAP-2*, or *fd59a* using knockin reporter alleles, combined with *fruitless* reporter expression. Generation of *Fer1-T2A-GAL4* is described in the Methods and shown in Extended Data Figure 3. White arrows: CREa1B axonal tract. Each gene labels CREa1B as well as a distinct pattern of other cerebral neuron bundles. For example, both *Fer1* and *dsf* are also expressed in the SMPad1A hemilineage. **f**-**h**, HCR RNA FISH or immunostaining for indicated transcription factors in CREa1B somata (outlined) at indicated timepoints. All factors are expressed consistently from at least 12h APF until adulthood. **i**, Two-photon images of the split-GAL4 intersections of hemilineage transcription factors, labeling CREa1B neurons. **j**, Model of hemilineage patterning downstream of the CREa1 neuroblast. Here and throughout, two-photon imaging of native fluorescence and confocal imaging of fixed and stained brains produce slightly different brain dimensions.

We used knock-in alleles in *TfAP-2*, *dsf*, and *fd59*a, and generated a T2A-GAL4 knock-in translational reporter of Fer1, to examine their expression over space and time (Fig. 2e, Extended Data Fig. 3). Across the brain, each marked a unique set of major tracts and clumps of neuronal somata, consistent with expression in diverse hemilineages and with previous description of “packets” of cerebral neurons sharing the same transcription factor^17^. We used these morphologies, combined with RNA FISH and immunostaining on the R19C05*^Pdfr^* lineage trace background, to demonstrate that only CREa1B expresses all four, confirming that our transcriptional cluster corresponds to this anatomic hemilineage (Fig. 2c, f-h’). Expression of all four endured from 12h APF to adulthood, similar to descriptions of hemilineage-specific transcription factors in the VNC (Fig. 2e, i)^35,36^. We also confirmed the pervasive expression of *dsf*, *Fer1* and *tup* in the SMPad1A hemilineage; and *Fer2* in the CREa1A hemilineage, and matched these to transcriptional clusters (Extended Data Fig. 4,5,6). These results support that the combination of our labeling strategy and our bioinformatic approach drives cells into their hemilineages.

Previous case studies have identified instances of both concordance and discordance between expression of a transcription factor in a neuroblast and in its differentiated daughters^30,37^. To ask whether the CREa1 neuroblast had expressed the TFs we observed in CREa1B neurons, we drove lineage tracing using *Fer1*, *TfAP-2*, and *dsf*. *Fer1* didn’t label any lineages using this strategy; *TfAP-2* labeled many lineages, but not CREa1; and *dsf* labeled most or all lineages (Extended Data Fig. 3). Conversely, CREa1 and several other neuroblasts in the R19C05*^Pdfr^* set express *vnd*^18^, yet we observed only sparse *vnd* expression in neurons (Extended Data Fig. 3). While these results do not exclude inheritance of neuroblast factors, they suggest a complex logical interaction between neuroblast spatial patterning and Notch signaling at neuronal birth to produce distinct TF codes in sister hemilineages of a single neuroblast.

### Complex transcription factor combinations uniquely define hemilineages

To test this model, we ascertained whether other transcription factors behaved “hemilineagely” in our dataset. In parallel with formal computational analysis of cluster markers (Methods), we blinded ourselves to TF identities and then manually inspected expression patterns of all 628 TFs across our cluster map split by developmental stage. We identified 89 TFs that had pervasive, enduring expression in subsets of clusters, of which 45 had a homeodomain fold (Fig. 3a, Extended Data Fig. 7). This three-fold enrichment corresponds to the well-documented role of homeodomain TFs in neuronal diversification^38–40^. Many overlapped with hemilineage-defining TFs in the VNC^35,36^ and/or are expressed in subsets of neuroblasts^41,42^. Transcriptional clusters expressed a median of 11 of these “hemilineage factors.” In addition to the hemilineage TFs, we describe the expression pattern of each TF in the genome, including TFs consistently expressed, varying across maturation, varying within each hemilineage cluster, and reflecting more than one developmental axis (Supplementary Table 3).

**Figure 3.**
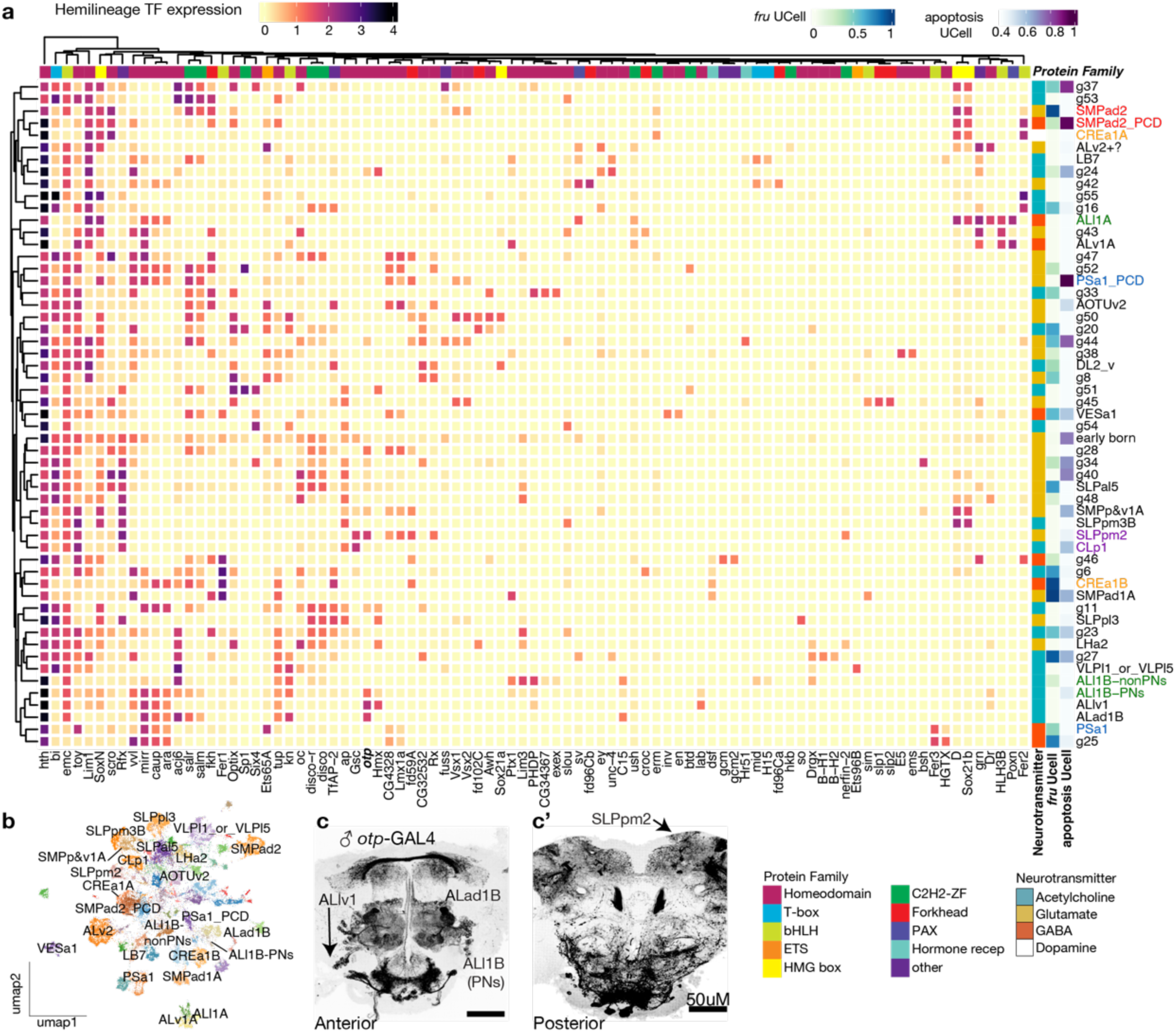
Complex transcription factor combinations uniquely define hemilineages. **a**, Heatmap displaying expression of 89 manually curated hemilineage TFs (columns) across integrated clusters (rows). Behavior of each *D. melanogaster* TF in our dataset is described in Supplementary Table 3. Colored pairs are sister hemilineages. We matched transcriptional clusters to anatomic hemilineages, and identified pairs of sister hemilineages, using data presented in Extended Data Figures 4 and 5, Supplementary Table 1, and the Methods. Transcription factors were partitioned into protein families using domains annotated in FlyBase or DAVID. Dominant neurotransmitter program for each cluster (based on *DAT*, *VGlut*, *VAChT* and *Gad1* transcripts) and UCell gene set enrichment scores for apoptotic programs (as described in Figure 1) and *fru* are displayed at right. *fruitless* expression is not linked to any single hemilineage TF. Rows and columns clustered by K-means. PCD: Clusters undergoing <u>p</u>rogrammed <u>c</u>ell <u>d</u>eath, which derive predominantly from p35 libraries in which we blocked the completion of apoptosis. A single cluster (“early-born”) contains primary cells of mixed hemilineage, based on *Imp* and *lov* expression shown below. **b**, Clusters in the integrated dataset that we matched to morphological hemilineages. **c**. Knockin reporter expression of an example transcription factor, *otp*, that labels a collection of hemilineage bundles. ALlv1, ALad1B, and SLPpm2 are included in the R19C05*^Pdfr^*-labeled hemilineages and identified by their unique tract morphology; *otp* expression is also visible in additional hemilineages that are not part of our transcriptional dataset.

To test the model that anatomic/developmental hemilineages are defined by a combinatorial “hemilineage transcription factor code,” we examined expression of 15 putative hemilineage TFs in the adult brain, often on the background of R19C05*^Pdfr^* lineage labeling (Extended Data Fig. 5 and 6, Supplementary Table 1). These factors were clonally restricted, expressed in hemilineage soma bundles, and their expression extended to adulthood. Combining our histology with prior literature, we matched the majority of frequently labeled R19C05*^Pdfr^* hemilineages to transcriptional clusters, including some that otherwise apoptose (Methods, Extended Data Fig. 5 and 6, Supplementary Table 1). In addition to CREa1, we were able to match the two living, sister hemilineages of the ALl1 and CLP1/SLPpm2 neuroblasts; as well as matching the SMPad2 and PSa1 natural hemilineages to their resurrected sister hemilineages. The TF codes for CREa1, PSa1, and ALl1 sister hemilineages had little or nothing in common, while those for CLP1/SLPpm2 and SMPad2 had some overlap (Fig. 3a).

### An orthogonal postmitotic TF code associated with neuronal birth order

In our analyses of TF variance across clusters, we noticed TFs that were expressed in swathes of neurons *within* each hemilineage cluster, including at 48h APF when neurogenesis has long ceased. We also identified a single cluster, 4, enriched for early-born cells expressing the temporal patterning RNA binding protein *Imp* and including neurons expressing early neuroblast temporal factors *Kr* and *pdm2*^13,24^. Combining “swath” TFs with those from cluster 4 yielded 36 TFs which we hypothesize vary downstream of neuronal birth order. Like the hemilineage TFs, this set was enriched for homeodomain proteins compared to the genome overall. In addition, many putative birth order TFs contain BTB (*ab, br, mamo, chinmo*) or pipsqueak (*dan, danr, Eip93F*) domains, or both (*CG3726*, *bab1*, *bab2, lov*) (Extended Data Fig. 7, Methods). Many of these 36 have been shown to act as temporal factors in neuroblasts; to label birth order cohorts within VNC or central brain lineages; or to allocate postmitotic neurons to alternative fates^13,21,24,43^. We confirmed by histology that Br, Pdm3, Mamo, and *lov* were expressed in subsets of neurons within hemilineages throughout the cerebrum (Fig. 4c, Extended Data Fig. 7). These factors showed complex patterns of transcriptional correlation and varied modestly across maturation stage (Fig. 4d,e)^58^.

**Figure 4.**
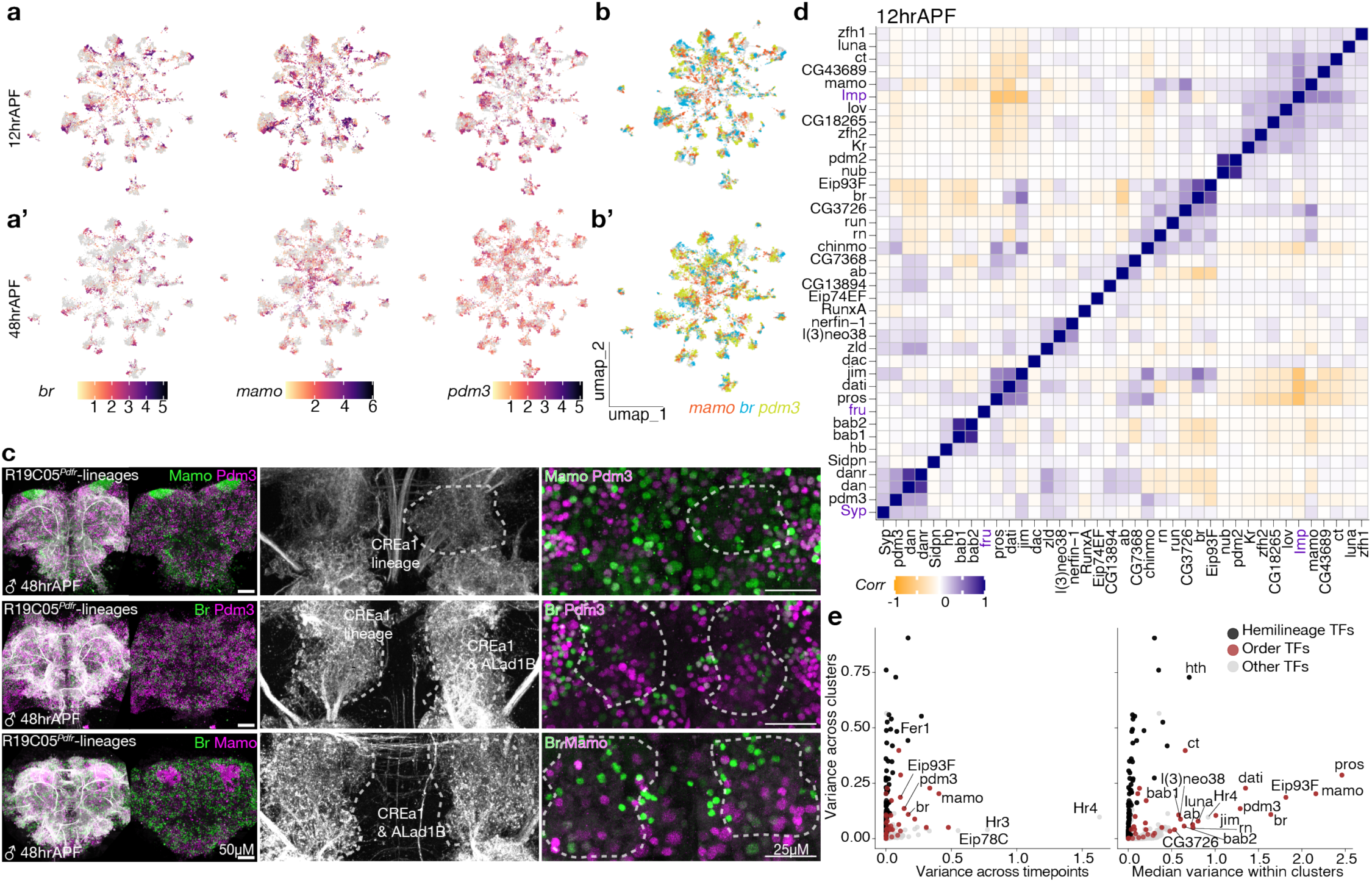
An orthogonal postmitotic TF code associated with neuronal birth order. **a**, **a’**, Featureplots showing expression of putative birth order factors *pdm3*, *br*, and *mamo* within hemilineage clusters. **b**, **b’**, Neurons within each cluster express *pdm3*, *br*, and/or *mamo*. **c**, Confocal images of R19C05*^Pdfr^* neuroblast clones combined with immunostaining for these putative birth order TFs. These proteins are largely non-overlapping within hemilineages (CREa1, dashed outline) and their representation in subsets of neurons is nearly universal across hemilineages. Bright patches of Mamo signal correspond to Kenyon cells, which are not from Type 1 lineages. **d**, Correlation of expression of 36 putative birth order TFs (manually curated, as shown in Supplementary Table 3 and described in the Methods). Correlation was performed across all cells from the 12h APF libraries. *Imp*, *Syp*, and *fru* are included for comparison. Putative birth order factors show complex patterns of correlation and anti-correlation with one another. *fruitless* is not linked to any specific birth order TF. **e**, Manually curated birth order TFs vary within clusters, and hemilineage TFs vary across clusters. Both sets are more stable across developmental timepoints than are previously identified maturation TFs (e.g. Hr3, Hr4)^58^. Factors that vary along more than one developmental axis are shown in Extended Data Figure 7 and described in Supplemental Table 3 and the Methods.

### Hemilineage TFs are required for CREa1B morphogenesis

Neurons in the same hemilineage share anatomic ground plan and excitatory/inhibitory valence, and this developmental axis is thus the major determinant of information routing in motivated behavior circuits. We hypothesize that hemilineage-specific TFs are required to give neurons within hemilineages their shared characteristics. We examined CREa1B in null mutants of *fd59a* and *dsf*, which survive to adulthood, and used the R19C05*^Pdfr^* enhancer to generate CREa1 clones with *TfAP-2* knocked down by RNAi. In all three cases, neurons were still specified and retained GABA immunoreactivity, but had clear morphological disruptions: Removing *fd59A* truncated axonal and dendritic projections, and we could not identify typical outgrowth tracts in *TfAP-2* knockdown clones (Fig. 5a-c, Extended Data Fig. 8). In *dsf* mutants, CREa1B neurite tracts are intact, but dendritic and axonal anatomy are disrupted (Extended Data Fig. 8). In each case morphological disruptions are shared by *fru* and non-*fru* neurons, suggesting these factors are required for the development of gross hemilineage features across subtypes. For *TfAP-2*, the effects we observe must be cell autonomous in postmitotic neurons, as we knocked it down clonally and found it was not expressed in the CREa1 progenitor (Extended Data Fig. 2, 3).

**Figure 5.**
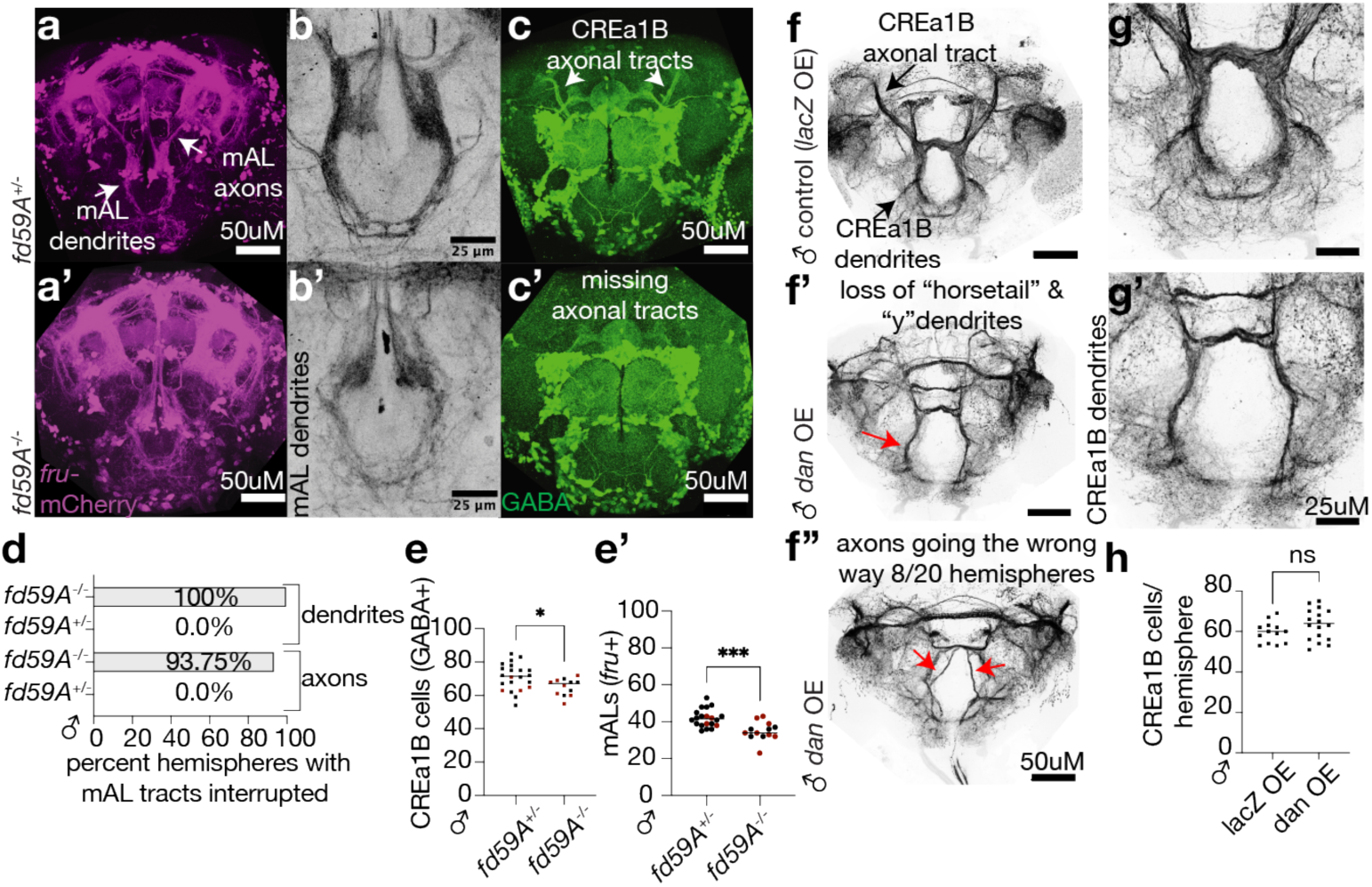
Proper expression of hemilineage and birth order TFs is required for neuronal development. Samples analyzed throughout this figure are male; female samples and additional analyses can be found in Extended Data Figure 8. **a-c**, Morphology of mAL neurons labeled by *fru*-mCherry or GABA immunostaining in controls versus *fd59a* mutants. (**a**, **c**) whole brain views; (**b)** displays mAL neuron dendrites zoomed in. Arrows, mAL axons or dendrites. **d**. Summary of the qualitative effect of the loss of *fd59A* on mAL axonal and dendritic morphology. n is shown by the number of points in (**e**). **e**, **e’**. Number of CREa1B neurons (GABA+) or mALs (*fruitless*-expressing) per hemisphere. Throughout this figure, significance: unpaired t-test; *: p<0.05, **: p<0.01, ***: p<0.001, ****: p<0.0001. **f-h**, Morphology and quantification of CREa1B neurons, labeled by *dsf*∩*TfAP-2* splitGAL4, in control and when the putative birth order factor *dan* is overexpressed. Y-shaped dendrites are lost in the overexpression condition (**g** versus **g’**), and axons are sometimes misrouted (as in **f”**). Extended Data Figure 8 shows phenotypes for additional hemilineage and birth order TF manipulations.

Loss of each TF impacted the number of cells in the hemilineage and the distribution of *fru* mAL neurons versus non-*fru* hemilineage cousins (Fig. 5e, Extended Data Fig. 8). Consistent with our previous identification of cell-type-specific *fru* regulatory programs^44^, no single hemilineage or birth order factor was predictive of *fru* expression across our dataset. Nevertheless, clonal knockdown of *TfAP-2* strongly reduced the number of *fru*-expressing CREa1B neurons in both sexes (Extended Data Fig. 8).

### Putative birth order TFs shift subtype patterning

Neurons with different birth order from the same hemilineage traverse the same tract highways but diverge in fine axonal and dendritic anatomies. CREa1B anatomies have been birth-ordered using tsMARCM in females^18^, which showed that diverse single-copy neurons with complex dendritic “wings” are born first; then neurons with a y-shaped dendrite are generated over a long larval span; and neurons with a simple dendrite and “antler” axon are born last. In addition, male mAL neurons have a straight “horsetail” dendrite, though these have been only roughly birthdated^26^. We constructed a splitGAL4 that labels all these subtypes by intersecting the *dsf* and *TfAP-2* hemilineage TFs, and used it both to label the hemilineage and to drive ectopic expression of putative birth order TFs. (Using this splitGAL4 to drive neuroblast lineage tracing did not produce any labeling, thus ectopic expression is limited to postmitotic neurons.) Ectopic expression of Pdm3 or Dan each shifted subtype distributions: When Pdm3 expression was induced throughout the hemilineage, female *fru* (i.e. mAL) neurons were lost, Br-expressing neurons were reduced in number, and the antler morphology characteristic of the late-born fate was gained at the expense of the wing morphology characteristic of the early-born fate (Extended Data Fig. 8). When Dan expression was induced, males lost the horsetail mAL morphology, and both sexes lost the y-shaped dendritic type; there was a gain in the early-born winged morphology (Fig. 5f-g, Extended Data Fig. 8). These results confirm that putative birth order factors, when expressed in postmitotic neurons, can causally alter the development of birth-order-specific morphologies, with Dan and Pdm3 induction mimicking earlier- and later-born morphologies, respectively.

### Birth order and sex diversify CREa1B transcription

Finally, we sought to understand how *fruitless* mAL neurons arise from within the CREa1B hemilineage. In females, the winged, y-shaped, and antler CREa1B morphologies described above can be resolved into 8 subtypes^18^; male-specific types and those that typically apoptose add additional complexity. We re-clustered the CREa1B set of cells and resolved 12 transcriptional subclusters, consistent with the expected diversity. mAL neurons are sexually differentiated due to differential splicing of the *fruitless* transcript, which generates functional FruM protein only in XY cells: Of 30-40 *fru* mAL neurons specified per hemisphere, 80% apoptose in female^26^; male mAL cells, which make FruM protein, have the horsetail dendrite and straight axon, while the surviving female *fru* mAL neurons have a y-shaped dendrite and ring-shaped axon^26^. We identified two *fru*-expressing regions of the cluster map, which were the only regions where male and female cell transcriptomes diverged (Fig. 6a). Comparing these two regions, we found they were differentiated by birth order factors—prominently *pdm3* and *br*—as well as by expression of pro-apoptotic genes among female cells of the *pdm3* set. (Those female cells derived from libraries in which we blocked apoptosis) (Fig. 6b). We therefore hypothesized that mAL*^pdm3^*cells are those programmed to apoptose in female, while mAL*^br^*cells survive. We confirmed co-expression of *br* and *pdm3* with *fru* by histology (Fig. 6e).

**Figure 6.**
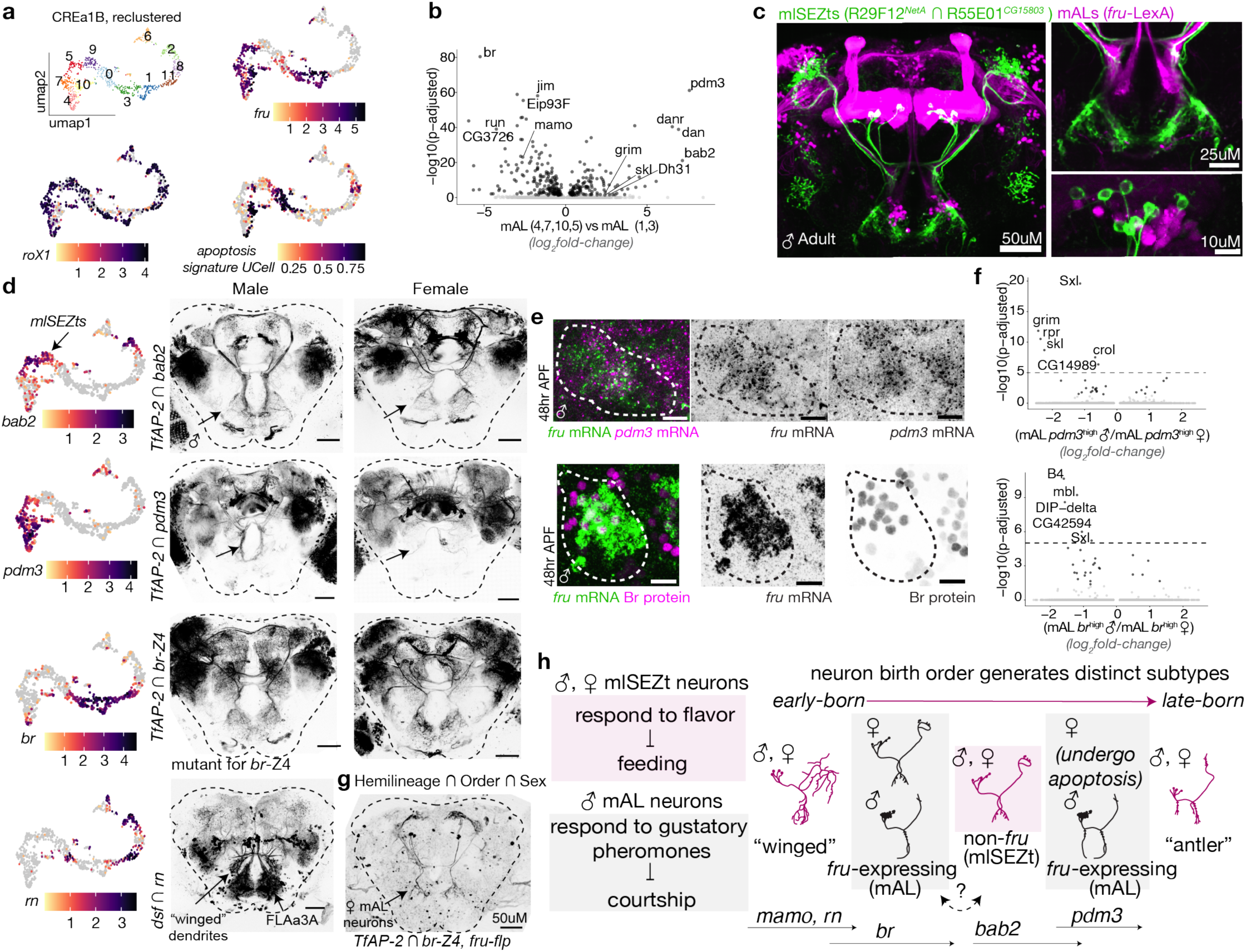
Birth order TFs differentiate mAL neurons and sex-shared sisters. **a**, UMAP of subclustered CREa1B neurons, integrated across timepoints. Featureplots display log-normalized expression. Subclustering was performed using all variable features. **b**, Volcano plot showing differential gene expression analysis between the two *fruitless* domains of the UMAP in (**a**). Black dots, significant based on Wilcoxon Rank Sum test. Gene labels highlight putative birth order factors and cell death genes, as described in the text. **c**, Comparison of mlSEZt and mAL neurons. At left, maximum intensity projection of confocal stack, at right, substacks focused on dendrites or somata. mlSEZt somata lack *fru*. **d**, Maximum intensity Z projections of splitGAL4 intersections of CREa1B hemilineage TFs and putative birth order TFs, as indicated. Arrows: CREa1B dendrites. Some morphologies in Z projections are obscured by neurons of other hemilineages, like FLAa3 (labeled). **e**, Histology for *fru* RNA with *pdm3* RNA or Br protein. Dashed outline: CREa1B cells. **f**, Volcano plots showing differential gene expression analysis between male versus female *pdm3* or *Br* mAL neurons. Top: Black dots, significant based on Wilcoxon Rank Sum test. Labels: protein coding genes, -log10(p-adjusted) > 5 based on Wilcoxon Rank Sum test. Bottom: Black dots and labeled genes, significant based on Wilcoxon Rank Sum test. *Sxl* is a member of the sex differentiation hierarchy. **g**, Model displaying circuit roles and likely birth order of CREa1B subtypes, and expression of order TFs and *fruitless*. Whether the two *fruitless*/mAL domains are contiguous remains ambiguous; the ordering shown is parsimonious with a single window of *bab2* expression and ordering of transcriptional types along the UMAP.

We searched the literature for neurons sharing the CREa1B outgrowth tract. Remarkably, we identified mlSEZt neurons, which have an analogous function to mAL’s, but participate in a different circuit: Where sex-differentiated mAL neurons gate male mating in response to pheromone tastes, mlSEZt’s are sex-shared and gate feeding in response to flavors^45^. We confirmed that mlSEZt somata are intermingled with mAL’s but lack *fruitless* (Fig. 6c). Strikingly, the mlSEZt neurons of both sexes have identical morphology to female mAL neurons (y-shaped dendrite and ring-shaped axon), despite their molecular discriminability via the mlSEZt splitGAL4 and *fruitless* (Fig. 6c).

To link additional CREa1B subtypes to birth order TFs, we intersected *bab2*, *pdm3*, *br*, and *rn* with CREa1B hemilineage TFs, using the splitGAL4 approach; we note that as *br* is on X, males with this knock-in allele are mutant for the *br-Z4* isoform (Fig. 6d). We found that *bab2* labeled both horsetail mAL neurons in male and the y-shaped morphology, which in male is represented only by mlSEZt neurons. *rn* labeled neurons with elaborate dendritic wings, consistent with its expression in early-born cells. *pdm3* labeled male mAL neurons, which were absent in female, confirming that this is the set that undergoes programmed cell death unless rescued by FruM. In females, *br* labeled neurons with the shared mlSEZt/female mAL morphology. We confirmed genetically that these include *fru* neurons, and thus that mAL*^br^*cells indeed survive in females. Males with this *br* allele had feminized mAL morphology and lacked *fru*, consistent with a requirement for Br-Z4 to allow *fru* expression in this window. Together with the birth-ordering previously performed^18^ and our genetic manipulations (Fig. 5f-g, Extended Data Fig. 8), these results confirm that putative birth order TFs are indeed linked to production of the specific morphologies generated over developmental time.

Blocking cell death in CREa1B equalized the proportion of mAL*^br^*versus mAL*^pdm3^* types between the sexes and allowed us to compare other aspects of their transcription (Extended Data Fig. 9, Fig. 6f). The differences in transcription between birth-order subtypes within a hemilineage (i.e. mlSEZt, versus mAL*^pdm3^*, versus mAL*^br^*) were greater than differences between sexes (e.g. mAL*^br^* male versus female)(Extended Data Fig. 9, Fig. 6f, Supplementary table 4). This suggests that FruM tweaks a sex-shared cell type that arises from the interaction of hemilineage and birth order patterning, rather than producing a completely different cell type^31^. In mAL*^pdm3^*’s, FruM’s most prominent impact was to suppress pro-apoptotic transcripts *grim*, *rpr*, and *skl*. Loss of mAL*^pdm3^*’s to cell death also removed expression of the neuropeptide *Dh31* in the female CREa1B hemilineage, as this peptide was restricted to the *pdm3* temporal cohort (Extended Data Fig. 9). While mAL*^br^* cells survive in both sexes and are sexually differentiated, transcriptional differences were modest: ∼20 genes--including a neuronal adhesion molecule (*DIP-delta),* a neuropeptide *(Nplp1)* and an RNA-binding protein *(mbl)*--were downregulated in male mAL*^br^*’s, consistent with direct repression by FruM (Fig. 6f, Extended Data Fig. 9, Supplementary table 4)^44^. Thus, sex differentiation occurs by conservative transcriptional adjustments within the plan laid down by hemilineage and birth order, supplying a male-specific variant to the male-specific mating circuit.

## Discussion

Here we describe a model for the diversification of rare neuronal types with unique behavioral roles. While motivated behaviors are well studied at the circuit and behavioral level, the transcriptional logic generating these immensely diverse cell types has been obscure. By capturing and sequencing postmitotic neurons of specific Type I lineages of the arthropod cerebrum, we identified two orthogonal TF codes, one of which diversifies hemilineages, and the other the neurons within hemilineages. Hemilineage tract morphology is derived from neuronal origins and is recognizable in connectomes and across individuals and species^8–10,15–17^. We show that this dominant organizational feature of cerebral circuits is produced by dedicated transcriptional programs, such that hemilineage is a Rosetta Stone linking ontogeny to molecular, anatomic, and functional axes. While neurons of the same hemilineage share gross projection pattern and neurotransmitter, subtypes diversified by birth order transcription factors have diversified axonal and dendritic morphologies and can thus supply the same computational module to distinct circuits^18,21,46^. Hemilineage and birth order transcription factors thus serve as barcodes of identity both in a causal developmental sense and from our position as observers of the brain. Our model thus decomposes cerebral “terminal selectors” into two groups defined by the separable developmental axes that induce them^47^. Finally, both hemilineage and birth order factors set the table that sex differentiation acts on.

Coincident manuscripts preprinted during the review period reached similar models of the roles of lineage, birth order, and sex in transcriptional differentiation of Type I lineages^48–50^.

### Paths from lineage to transcription factor expression

Transcription factors are re-used in different neurogenesis programs: Many of the birth order factors we describe pattern VNC and/or Kenyon cell temporal cohorts, and many also act as temporal factors in VNC or cerebral neuroblasts^13,24^. Members of our hemilineage transcription factor set participate in multiple combinations in the cerebrum, and likely in additional combinations within VNC hemilineages^35,36^. Many in the set also serve as neuroblast spatial factors and as temporal or spatial factors in the visual system^13,41,51,52^. The re-use of these factors across different neurogenic events likely relies on a diversity of upstream paths through which they can be switched on^53^. Similarly, we found that *fruitless* expression emerged out of diverse hemilineage and birth order contexts and depended on subtype-specific upstream factors.

While singular cerebral neuroblasts can be identified based on differential transcription factor expression^41,42^, few have been matched to the anatomic hemilineages they produce; the TFs we test here are disjoint between the neuroblast and its differentiated daughters, suggesting that linking neuroblast transcriptional profiles to those of their daughters may be complex. In a few cases, handoffs have been observed between neuroblast spatial factors and those in differentiating daughters^54,55^, raising the potential for a web of regulatory interactions guiding the emergence of hemilineage TF combinations downstream of spatial and Notch developmental axes.

### Diversification downstream of hemilineage and birth order TFs

Spatial versus temporal patterning dominate neuronal diversification in different types of circuits: Sister neurons of cerebral or nerve cord hemilineages supply a common computational module to distinct circuits, with birth order factors working within the hemilineage groundplan. Spatial patterning is similarly prominent in the vertebrate spinal cord, which also relies on Notch and birth order diversification^56^. In contrast, the columnar organization of visual or cortical circuits results from temporally diversified daughters of the same progenitor wiring together. In these circuits, the degree of spatial diversification of neurons is lower, while the transcriptional and morphological differences produced across birth orders is higher. The degree of differentiation produced along specific developmental axes likely results from the characteristics of the TFs that vary along that axis, in particular whether they can activate gene expression patterns de novo or only edit regulatory events laid down by others^57^. In the arthropod cerebrum, we find that BTB and pipsqueak domain proteins, which often act as repressors, are enriched among, and specific to, the birth order TFs.

### Extant versus potential TF codes

We were struck by the apparent excess of TFs devoted to hemilineage patterning: The 89 postmitotic hemilineage TFs we identify, taken 10 at a time, could produce billions of unique combinations, yet there are <200 cerebral hemilineages. Thus, the TF combinations that exist in the brain are a very sparse subset of the combinations that could be formed from the individual factors. Recent analyses in the visual system have found that the amount of transcriptional diversity that could be produced by the available transcription factor set is also far in excess of the number of cell types actually made^52^. How these cocktails are generated, and what is special about those combinations that occur in extant brains, awaits further study.

## Acknowledgments

We thank Tzumin Lee, Haluk Lacin, Yu-Chieh David Chen, M. Neşet Özel, Ted Erclik, Mubarak Syed, Jim Skeath, and Uwe Walldorf for developing and sharing critical fly alleles, composite genotypes and antibodies, without which this work would not have been possible. We also thank the Bloomington Drosophila Stock Center. Margarita Brovkina provided conceptual and technical input. Veronica Sikora collected mlSEZt images. FAC-sorting was performed with Kamlai Saiya-Cork in the BRCF Flow Cytometry Core. Library prep and next-generation sequencing was carried out in the Advanced Genomics Core at the University of Michigan. Research reported in this publication was supported by the University of Michigan Advanced Genomics Core, the UM Single Cell Spatial Analysis Program and the National Cancer Institutes of Health under Award Number P30CA046592 by the use of the following Cancer Center Shared Resource: Single Cell and Spatial Analysis Shared Resource. Funding was provided by NINDS DSPAN F99NS139459 and CMB T32 awards GM145470 and GM007315 to NE; NIDCD R01DC018032, McKnight Scholar Award, Pew Biomedical Scholar Award, and startup funds from the University of Michigan to EJC; and National Science Foundation (IOS-1845673) to TRS.

The authors declare no competing interests.

## Methods

### Fly husbandry

Flies were maintained on Bloomington food with a yeast sprinkle (“B” recipe, Lab Express, Ann Arbor, MI) at 25C on a 12:12 light-dark cycle with at least 60% humidity (provided by a beaker of water in the incubator). For staging developmental time points, pupae were marked at the 0hr APF “white pupae” stage. Animals for scRNAseq were collected +/−1-2 hours from the stated developmental time points (e.g. 48hr APF were collected from 47-49hr APF).

### Flow cytometry

Flow cytometry to sort GFP-positive neurons from pupal brains was performed similarly to adult brains^44^: We pretreated plasticware including tips and eppendorf tubes with PBS + 1% BSA to prevent cells from sticking to the plastic. Dissections of fly brains were done in Schneider’s+1% BSA for up to one hour. Optic lobes were removed for the 48hrAPF time points. After dissections, Collagenase was added to the brains at a concentration of 2mg/ml in Schneider’s+1% BSA then placed at 37C for 12 minutes (versus 20 minutes for adult brains). Afterwards, we did a manual dissociation by pipetting the brains up and down at least 30 times, then spun down the cells at 300g for 5 minutes. We removed the supernatant, then resuspended the pellet in 200ul of PBS + 0.1% BSA, passed it through a cell strainer cap, and spun it down briefly to collect cell suspension at the bottom of the tube. We added 50ng/mL of DAPI to our sample and then sorted them on a FACSAria III (University of Michigan BRCF Flow Cytometry Core) into a PBS + 1% BSA cushion. During flow cytometry, dead and dying cells were excluded using DAPI signal, and forward scatter and side scatter measurements were used to gate single cells. Scatter profiles consistent with somata (rather than neurite fragments) were determined by back-gating on DAPI. Using our dissociation methods, ∼70–90% of singlets appeared viable (DAPI-low) for both time points. Because the cells were to be analyzed using single cell sequencing, we set our gates generously (i.e. tolerated sorting false positive cells so as to capture all true positives). With these generous gates, 9-13% of these viable cells were gated as GFP+ for our 48hrAPF time points, and ∼2% for 12hrAPF. During sorting, we made two adjustments to protect the fly primary cells, which were very delicate—we disabled agitation of the sample tube, and sorted using the “large nozzle” (100μm, i.e. using larger droplet size and lower pressure).

### scRNA-seq quality control metrics

We prepared 8 libraries using the lineage labeling strategy: 2 replicates at 48hrAPF, 2 replicates at 48h APF with apoptosis blocked, 2 replicates at 12hrAPF and 2 replicates at 12h APF with apoptosis blocked. Potential doublets and low-quality cells were filtered out by discarding cells that didn’t express at least one read for any of our transgenes (lexAop-GFP, lexAop-p35 or LexA-p65), cells with >50000 UMIs, or cells with < 1000 unique features. The ∼10% of cells with the highest percentage of mitochondrial transcripts were discarded for all libraries. Features corresponded to genes that were expressed in at least one cell across libraries and were filtered for any mitochondrial genes, transposable elements, ribosomal proteins, mod(mdg4) trans-spliced precursors, and non-coding RNAs besides *lncRNA:roX1* and *lncRNA:roX2* (which are used to assign sex to cells). After filtering, we had a total of 57,076 cells and the number of features used for downstream analyses was 12,957.

### scRNA-seq clustering and integration of libraries

We merged all the libraries and used the Seurat package (Seurat V5 (Seurat_5.3.0))^79^ to cluster and integrate our datasets and to visualize our clusters across conditions (+/- p35) and timepoints (12 vs 48hrAPF). To do this, we normalized gene expression using the NormalizeData function (LogNormalize method). Rather than finding and using the most highly variable genes for the subsequent steps, we used a list of 628 transcription factors from FlyBase – 609 of which were expressed in our cells. We then scaled our data (ScaleData) while regressing out mitochondrial gene expression (percent.mito) and sequencing depth (nCount_RNA). We then ran principal component analysis using the scaled expression of our 609 transcription factors (RunPCA) otherwise using default parameters. For clustering and visualizing, we used default parameters, including 50 principal components based on an ElbowPlot (0-200) (FindClusters, FindNeighbors and RunUMAP). We then integrated our datasets (IntegrateLayers) using Harmony Integration and ran the clustering (FindClusters, FindNeighbors and RunUMAP) as we just described above to get an idea of the cell types and cell type diversity that existed in our dataset.

To enrich for non-MB Type I hemilineages, we removed Kenyon cells, glia and 12hrAPF-specific clusters we found to be optic lobe neurons or progenitor cells (Extended Data Fig. 1): In our top-level clustering, Cluster 5 contained Kenyon cells based on the expression of *Mef2*, *ey* and *dac.* Cluster 13 was glia based on the expression of *repo.* Cluster 3 primarily contained cells from a single library (low quality cells), so we removed it. 7 clusters (20, 15, 6, 22, 46, 27, 43) were specific to the 12hrAPF time point and were enriched for *scro* but depleted of *Imp* and *dati*^93^, indicating they are optic lobe neurons; we removed these. (At 48h APF, we removed optic lobes during our dissections; at 12h APF, the optic lobes have not yet separated from the central brain and are thus present during our sorting.) 21 and 47 expressed *dpn* or *dap* and were presumed to be progenitor cells and removed. Some of these cell types are truly labeled by our recombination-based strategy (progenitors, glia deriving from Type I lineages, sporadic clone induction in KC lineages). Others are false positives due to a combination of our generous gating and their prevalence. After filtering, we retained a total of 42,844 neurons (18,854 cells from 48hrAPF and 23,990 cells from 12hrAPF). Extended Data Figure 1 has the breakdown of how many cells come from each library. To generate the final clustering map we used for biological analyses, we re-ran the workflow above, except that after integrating the datasets, we used a resolution of 1.4 in FindClusters determined using ClusTree^81^. This produced relative stabilization of 56 clusters that we assign as postmitotic neurons originating from cerebral lineages. These are almost all Type I lineages, though we observed rare labeling of Type II sublineages as shown in Supplementary Table 1.

As we describe in the results text, we hypothesize that these clusters mainly corresponded to hemilineages, with larger clusters representing hemilineages always labeled by our lineage trace, and smaller clusters representing hemilineages that were labeled more rarely. That said, as we worked back and forth between the transcriptional to histological points of view, we observed that there was no single clustering resolution that perfectly reflected hemilineage relationships: At lower resolution, some clusters contained multiple hemilineages (e.g. anterodorsal and lateral olfactory projection neuron hemilineages), while at higher resolution, some hemilineages split into more than one cluster along the birth order axis. We chose a resolution that kept most hemilineages unified and manually split some of our clusters, as shown in (Extended Data Fig. 1i-j). For example, cluster 0 contained the *DAT*+ CREa1A neurons, which express *scro*, *dmrt99B*, *erm*, *D, Lim1*, *hth, SoxN, CG9932, Sox21B* and *Fer2*, along with other cells that strongly shared *Fer2, scro, D, Sox21b, hth* and *Lim1* but lacked DAT and expressed *Gad1* and appeared to be destined to apoptose based on expression of *grim*. This suggested they are a resurrected hemilineage that typically undergoes apoptosis, which we found experimentally was from SMPad2. We split this cluster into CREa1A and SMPad2_PCD. We also manually split cluster 11 and 41, which contained both Alad1B (adPNs) and a small set of ALl1B/lPNs based on the expression of *acj6* (Alad1B/adPNs) versus *vvl* (ALl1B /lPNs). The matching of transcriptional clusters to hemilineages is described in Supplementary table 1.

### Identifying the lineages labeled by R19C05^Pdfr^ and their hemilineage sisterships

To identify the lineages included in our set of clones, we collected 10 male and 10 female R19C05*^Pdfr^* lineage trace brains without p35 (“natural” lineages) and 10 male and 10 female brains with p35 (which include “resurrected” cells). We performed IHC against GFP lineage labeling together with neuropil stain with anti-Brp (nc82) to allow image registration. Brains were cleared by mounting in SlowFade Glass. We imaged stained brains at 0.38 micron XY resolution and 1 micron Z steps (as for most researchers studying the cerebrum, our Z dimension is anterior-posterior) using a 40x oil immersion objective, and registered them to the JRC2018 Unisex common brain template^94^ using non-rigid registration via CMTK. Because GFP in this strategy labels neuronal membranes, we could use both the positions of labeled soma and the morphologies of labeled tracts to identify hemilineages, which we did by comparing to the FAFB connectome, where hemilineages are annotated (see below). For the “natural” genotype, we took a census of each of the 40 hemispheres (Extended Data Fig. 2c) to determine the labeling rate for each lineage. Overall, we labeled 13 lineages in at least half of brains, and 13 more at least once but less than 20 times in our census. This genotype was originally described in Ren et al.^22^, who catalogued 10 labeled lineages. We found 8 of these in the set of 26 in our census. We observed a few additional lineages labeled sporadically in our histology experiments that we didn’t observe among the 26 in our census, and note that our 8 sequencing datasets incorporated about 400 total brains.

While a number of papers had studied cerebral neuron anatomies and origins previously, adult anatomic hemilineages were systematically named and catalogued in a set of 2013 papers from the Lee, Ito, and Hartenstein labs, and a 2010 census of the lineage origins of *fruitless* neurons provided additional clonal data for the cerebrum^3,8–10,95^. Two nomenclatures (Ito/Lee versus Hartenstein) were reconciled and anatomic hemilineages were annotated in the FAFB connectome by Volker Hartenstein and Alexander Bates (Alex Bates, personal communication)^15,96^. These annotations have been used to seed hemilineage matches in other connectomes^97,98^. Most of the annotations in the connectome set include only the secondary parts of hemilineages (i.e. the large sets of cells born in the larva); primary neurons, born in the embryo, are mainly categorized as “putative_primary,” and await association with the rest of their hemilineages. As many have noted, primary neurons share aspects of their anatomy with the secondary parts of their hemilineages, but often have unique and more “fancy” innervation patterns and larger somata^99^. Morphological subgroups from Type II neuroblasts are also typically referred to as “sublineages” rather than “hemilineages,” because (1) there are more than two major morphologies produced by Type II lineages, and (2) the lineage tree is structured differently for Type II lineages, involving two temporal axes (neuroblast and intermediate progenitor)^100,101^; in the connectome metadata, however, Type II sublineage anatomies are simply called “hemilineages.” (For neuroblasts such as ALl1 that produce more than one major morphology within a hemilineage, those different morphologies may also be referred to as “sublineages.”)

Because the two hemilineages from a single neuroblast are generated in parallel (i.e. each GMC division produces one neuron for each hemilineage), and because our lineage labeling here is restricted to initiation in Dpn+ cells, there is no way to label one Type I hemilineage without labeling its sister. (Dpn is also expressed in intermediate progenitors from Type II neuroblasts; while we find that R19C05*^Pdfr^*is mostly restricted to Type I neuroblasts, we occasionally label a Type II sublineage, as detailed in the supplemental table.) While hemilineages are clonally related and generated in parallel, the two sets of soma are usually pulled apart from one another as their primary neurites grow out along different tracts (this is why the cells expressing hemilineage-specific TFs, like *Fer1* in Fig. 1c, are not mingled with their TF-negative sisters); in a minority of cases, the two hemilineages radically separate during metamorphosis (this is why there is sometimes ambiguity in identifying sister hemilineages)^8–10,95^. A survey of 18 Type I neuroblasts suggested that ∼40% of cerebral Type I neuroblasts produce two living hemilineages, while 60% produce a “lone” living hemilineage whose sister hemilineage apoptoses^18^. The lineage tracing datasets from 2013 identified many cerebral hemilineage pairs that derive from the same neuroblast, and also provide clues of cases in which a sister is suspected but hasn’t yet been identified—lineages that produce more than one “cell body fiber,” (Ito), that always occur together (Lovick, Wong), or that produce two first-born neurons observed in the adult (Yu)^8–10,95^.

Using these clues together with our own clone set, we exhaustively examined whether specific “lone” hemilineages always occurred together, likely to be unidentified sister sets. CLP1 and SLPpm2 were a pair that we newly identified as sisters. In addition, we found cases in which a full living hemilineage is paired with a small set of very different neurons, likely primary neurons of the sister hemilineage (born in the embryo) for which the secondary part of the hemilineage (i.e. the large group of neurons born in the larva) apoptosed. PSa1 and SMPad2 are examples of this type of lineage. To find these “full hemilineage plus primary neurons of the second hemilineage” cases, we were aided by (1) the Cachero 2010 dataset, which used clonal labeling to identify *fruitless* cell types from different lineages; (2) descriptions of the central complex cell types and their origins; and (3) images of clones from these and other papers. For example, for PSa1, we saw that the lineage clone in Cachero 2010 for aDT5, which is from PSa1, also include aDT7 neurons^3,8^. In the Ito et al. PSa1 clone, these aDT7 neurons are also visible. In the connectome, aDT7 cells are annotated as “putative_primary,” and like most such “putative_primary” neurons, they are not assigned to a hemilineage. Unlike the annotated PSa1 hemilineage, which are predicted to be GABAergic, aDT7 cells are aminergic. In hemispheres where we label PSa1, we always label aDT7 cells. We can now assign them as sister to PSa1.

We also examined whether any of our hemilineages of interest in FAFB could be mixtures of two different hemilineages from the same lineage—we looked for cases where an annotated “hemilineage” included cells with different neurotransmitters that follow distinct outgrowth tracts (Alex Bates, personal communication). We took into account the number of cells annotated to each hemilineage across FAFB, BANC, and maleCNS^15,97,98^. We also used community annotations in Codex to connect well-studied neurons to their hemilineages.

Finally, we compared our census brains with and without p35, to look for cases of resurrected whole or partial hemilineages. While this seemed initially like it would be straightforward, we were humbled by the complexity of programmed cell death in brain development, which we understand can occur through at least four routes: First, one of the neurons from each GMC can be programmed to die as it is born, at embryonic or larval stages, as observed in Lin 2010^6^. Second, some neurons that functioned in the larval brain apoptose during metamorphosis^102^. Third, neurons can be removed by trophic cell death, i.e. when they are overproduced relative to their partners^103^; while we haven’t yet observed this in the cerebrum (despite tinkering extensively with ratios of pre-and post-synaptic cells in our work in the mushroom body in Elkahlah et al., 2020; Ahmed et al., 2023; and Pan et al, manuscript in preparation^104,105^), we don’t think it should be ruled out. Fourth, there is extensive sex-specific apoptosis in pupal stages^26^. Long story short, we were able to find some clear resurrection of whole hemilineages via ectopic tract morphologies (from the ALv1 and SMPpm1 lineages) and were able to identify the lineage origins of two scRNAseq clusters containing resurrected neurons (those from PSa1 and SMPad2). Cataloguing which cells apoptose from every lineage/hemilineage and the conditions under which they do so will require extensive further analysis and experimentation.

Clues from the literature and our data and reasoning for lineages in our set are provided in Supplementary Table 1. The references we used to identify lineages/hemilineages and relate them to each other are: ^3,6,8–10,15,18,20,30,95–98,106–113^. Overall, in our 26 lineages, we conclude that ¼ have two full living hemilineages; ¼ have one living full hemilineage and a “remnant” of primary cells from the other hemilineage; ¼ have one living full hemilineage and one apoptosing full hemilineage; and ¼ we did not have enough information to determine. Three of the anatomic sets we characterized include so many cells that they likely derive from more than one “twin” neuroblast, SMPad1, SMPad2, and “VLPl1 or VLPl5.” For the VLP groups, this is almost certainly the case, as we observe 1n and 2n quanta in different hemispheres.

### Matching anatomic hemilineages to transcriptional clusters

Once we had determined the ground truth of which lineages are labeled by our genetic strategy, we began the work of matching anatomic hemilineages to transcriptional scRNAseq clusters, guided by (and at the same time, testing) the hypothesis that we had clustered neurons in our dataset by hemilineage. We used many types of clues: Papers in the literature that had studied TF expression in the pupal or adult brain; markers of neurons that are constituents of these hemilineages, especially *fruitless* neurons, which are often the best studied representatives of their hemilineage; experimentally determined or EM-predicted neurotransmitters and rule of thumb that most hemilineages share neurotransmitter usage; patterns of apoptosis in the brain and expression of pro-apoptotic genes in transcriptional space; the expected stoichiometry of different lineages in our dataset; expectation that sister hemilineages would have similar stoichiometry; and a large set of histology experiments performed in this paper examining hemilineage transcription factors on the background of R19C05*^Pdfr^*lineage tracing. We performed these latter experiments in the adult as we are not yet expert enough anatomists to make hemilineage matches during development. This also confirmed that the hemilineage transcription factors we identified at 12 and 48h APF continue to be expressed in similar patterns in adulthood.

The evidence and reasoning for every match are provided in Supplementary table 1. We note that almost all the evidence we amassed was in concordance, and we eventually found explanations for most pieces of data that initially appeared to conflict. Ultimately the process was like the children’s logic puzzle in which one is asked to put characters in order around a table, given clues such as, “Sophie is sitting next to someone wearing a hat, Alfred doesn’t eat vegetables, the child with the hat is eating broccoli, one person is eating spaghetti, and there are no dogs sitting at the dinner table.”

The references we used to learn what cell types are included in hemilineages of interest and to make transcription-to-anatomy matches were as follows: *fru* or *dsx* cell memberships within lineages, or their molecular markers^3,4,10,22,31,113–117^. Other cell type memberships and molecular markers, by lineage: ALad1^35,118–122^; CLP1&SLPpm2^68^; CREa1^26,32,33^; VESa1^30^; ALv2^14,110,123,124^; PSa1^3,116^; LB7^125^; SLPpl3^17^; VLPl1_or_VLPl5^113,114^; SMPad1^32^; ALlv1^125,126^; ALv1^118,121,127,128^; AOTUv2^129^; ALl1^35,106,109,112,119–122,127,128,130^; LHl2 and LHl4^131^.

### Subclustering, integration and analyses of CREa1B cells

Subclustering was performed using the standard Seurat workflow: gene counts were normalized (function: NormalizeData); 2000 highly-variable genes (FindVariableFeatures) were scaled and used for PCA (functions: ScaleData/RunPCA); first 50 principle components were used for clustering (functions: FindNeighbors/FindClusters) and UMAP projections (function: RunUMAP). For integration we used the standard integration workflow of Seurat 5 using Anchor-based CCA integration to identify shared cell types across time (12hr and 48hr APF). It works by correcting the previously identified PCA coordinates and these corrected coordinates are for clustering (functions: FindNeighbors/FindClusters, resolution = 0.8, clusters = 11) and UMAP projections (function: RunUMAP) (Hao et al. 2024). Clusters 3 and 4 lacked *fd59A* and *TfAP-2*. Those cells were removed and the workflow above was re-run (955 cells, resolution = 1.2, clusters = 12). Resolution was determined using Clustree.

Proportion of mAL types in males vs females was calculated by dividing the number of *pdm3* or *br* expressing mALs for each sex by the total number of *fru* neurons for each condition (+/-p35).

For the comparison of wildtype female mAL neurons and mAL neurons that undergo programmed cell death. First, female mAL neurons were subsetted and then neurons with a high apoptosis signature determined by UCell using *grim*, *rpr* and *skl* (>=0.04) were compared to mAL neurons with a low apoptosis signature (< 0.04) using Seurat FindMarkers.

### Differentially expressed gene analyses

All differentially expressed gene analyses were done using the FindMarkers function using default parameters. Afterwards the differentially genes were filtered for any non-coding RNAs besides *lncRNA:roX1* and *lncRNA:roX2*, mitochondrial genes, transposable elements, ribosomal proteins and mod(mdg4) trans-spliced precursors. Plots were made using the ggplot2 package. CREa1B, CREa1A, cluster 0 (SMPad2_PCD) ALad1B (adPNs) and ALl1B (lPNs) marker genes were determined by comparing each of those clusters to all other clusters in the dataset (Supplementary table 2). In the case of CREa1B, clusters were split by time point first, then marker genes were found as described above (Supplementary Table 2).

### Categorizing transcription factor expression

#### Computational approach/ FindAllMarkers

For the differential gene expression analyses, we used the FindAllMarkers function from Seurat^79^, using the default Wilcoxon Rank Sum test on log-normalized feature counts for our 55 clusters (hemilineage annotated clusters). We selected for transcription factors that had a minimum percentage difference (min.diff.pct) between the two populations of 50% because we were looking for positive marker transcription factors for our clusters, resulting in 101 transcription factors. The results of this analysis are included in Supplemental table 1 (DEGs (TFs_0.5)).

### Categorizing transcription factor expression

#### Manual categorization of transcription factor expression

For manual categorization of transcription factor expression, we used Seurat “featureplots” to display the expression of the 609 FlyBase TFs that were expressed in at least one cell in our integrated dataset; we split our dataset by timepoint in order to discern maturation-based changes in TF expression. We blinded ourselves to gene names, and then categorized each expression pattern as “lineage” (expression in one or more, but not all, whole clusters), “order” or “order-like” (expression in subsets of cells within each cluster), or “possibly maturation” (broad expression, varying across time points). Transcription factors that did not fit into these three simple categories were categorized as either “consistent” (anything that was not differentiated across or within clusters, including near absence of expression, scattered positive cells, or pan-neuronal expression), “lineage-confusing,” or “unclassified/confusing” (usually a combination of the other axes), “Clk-related,” for circadian TFs co-expressed in a small Clk-expressing cluster, “Dsx” for doublesex, “fru” for fruitless, and rarely, multiple labels. Overall, we found 112 putative hemilineage transcription factors, of which 89 had cluster-defining and cluster-specific expression without varying over other axes. We identified 35 transcription factors we hypothesize reflect birth order. Our manual descriptions of the expression patterns of each TF are provided in Supplementary table 3. We manually added *chinmo* to that list of TFs, due to its known roles in temporal patterning and after re-inspection; it has a gradient, rather than binary, expression, as described in the literature^132^. This resulted in 36 TFs we refer to as “birth order TFs”. We focused our spatial TF analyses on the 89 TFs with simple cluster-defining expression, as described above, and we refer to them as “hemilineage TFs”. We note one cluster in which other aspects of patterning won out over hemilineage: Cluster 4, which is enriched for cells expressing *Imp*, *lov*, *nub*, *pdm2*, and *Kr.* Most of these are well-studied early neuroblast temporal factors^133,134^, suggesting that this is a cluster of primary cells of mixed lineage; additional cells expressing *Imp* and *lov* joined typical hemilineage clusters. To confirm that cluster 4 was not a hemilineage, we performed RNA FISH for *lov* on the R19C05*^Pdfr^* lineage trace background and found that within type I cerebral hemilineages, it labels a subset of cells per clone, not large bundles of soma (Extended Data Fig. 7c-d). Among the 101 computationally identified TF markers 82 of those TFs were contained in our list of 89 hemilineage TFs. 10 were called as lineage-weird and the rest were *fru*, *vri* (*clk*-related) or birth order TFs. The hemilineage codes in Supplementary Table 1 were determined by filtering the differentially expressed genes identified by Seurat (Supplementary table 2, (all DEGs)) for our list of manually identified hemilineage transcription factors.

### Lineage variability and temporal dynamicity scores

We also computationally assigned each gene in our dataset a lineage variability score and a temporal dynamicity score (Extended Data Fig. 6u). These scores were calculated using code generously shared by Saumya Jain and implemented similarly as in (Jain et al. 2022)^58^: We converted our cells into pseudo-bulk RNA-seq datasets based on their cluster identity and time (cluster 1 12hrAPF, cluster 1 48hrAPF…etc) by aggregating the expression of counts according to cluster and time (AggregateExpression)^79^ The lineage variability score was calculated by determining the variance in gene expression across lineages within either time point where the gene was expressed, taking the maximum of those two scores, and then normalizing by the average expression across cell types at the relevant time point. The temporal dynamicity score was calculated by taking the absolute difference between our 12hr and 48hr timepoints for each of our hemilineages/ clusters where the gene was expressed, and the averaged value across clusters was then normalized by the average of the peak expression. For a gene to be considered expressed it had to have an aggregated expression level of > 0.2. Both scores had their mean centered to 0 and were transformed to the natural log scale for visualization.

### Heatmap and other plots

Heatmaps were made using the ComplexHeatmap package in R in figures 3^88^.

We identified protein domains enriched within the 89 hemilineage TFs and 36 birth order TFs using DAVID Bioinformatics Functional Annotation Tool and FlyBase^135–137^.

The plots in Figure 4b-b’ were made by subsetting neurons that express (expression level > 0) either *br*, *pdm3* or *mamo* but not the other two (expression level == 0).

We used the UCell package^82^ to calculate the gene set enrichment scores at single-cell level of the following genes: 1. *VAChT*, *DAT*, *VGlut*, *Gad1*, 2. *fru* 3. *rpr, skl* and *grim* to assign our pseudobulk clusters a neurotransmitter identity, whether it’s sexually-dimorphic based on *fru* expression, and to an apoptosis score. The ranking in UCell uses the Mann-Whitney U statistic.

### Variance across vs within clusters vs across time

We calculated the variance in expression for each transcription factor across our pseudobulk clusters grouped by either cluster or time. Median cluster variance was calculated by finding the variance in expression for each transcription factor for each cluster and taking the median value for each transcription factor.

### fruMiMIC[mcherry] (fru-mCherry)

The *fru*MiMIC[mcherry] flies were made via recombinase-mediated cassette exchange (RMCE) performed by Rainbow Transgenic Flies Inc. with the pBS-KS-attB1-2-GT-SA-mcherry-SV40 vector ordered from the Drosophila Genomics Resource Center (DGRC Stock Number: 1324)^138^.The DNA was prepared with a Qiagen Plasmid Midi Kit and injected into the original *fru*[MI05459] line (BDSC: 42086)^60^. Transformants were crossed to yw double balancer flies and were screened for loss of y+ over the third balancer. As the swap-in allele can incorporate in either direction, we screened F1 progeny of animals positive for loss of the original MiMIC allele for *fru*-mCherry expression using our 2P microscope. (mCherry has a poor 2p excitation profile, but we captured the edge of its 1p excitation using ∼700nm illumination.)

### Fer1-T2A-Gal4

The Fer1-T2A-GAL4 construct was made using CRISPR/Cas9 HDR as previously described in (Xiao, Yuan, and Yadlapalli 2022)^139^. The donor cassette was designed so as to have ∼1kb homology arms flanking the stop codon (5’ PAM silently mutated, 3’ PAM deleted) with the T2A-Gal4 sequence, taken from addgene#125211^140^, immediately preceding the stop codon. The donor construct was synthesized and cloned into pBluescript II SK(-) by GENEWIZ from Azenta Life Sciences. Two Fer1 gRNA sequences were generated, as listed in the Oligonucleotides section, and each was synthesized as overlapping primers and annealed to generate dsDNA with BbsI overhangs. Each guide (5’ and 3’) was cloned into pU6-BbsI-chiRNA, generating 5’ and 3’ Fer1 gRNA plasmids. These two plasmids, together with the homology-directed repair donor, were cleaned and injected into *y,sc,v; nos-Cas9; +/+*^141^ by Rainbow Transgenic Flies Inc. Injected flies were crossed to double balancers and their progeny were screened by PCR genotyping of amputated legs using the primers listed in the Oligonucleotides section.

### Brain dissection and two photon imaging

Brains were dissected in external saline (108 mM NaCl, 5 mM KCl, 2 mM CaCl2, 8.2 mM MgCl2, 4 mM NaHCO3, 1 mM NaH2PO4, 5 mM trehalose, 10 mM sucrose, 5 mM HEPES pH7.5, osmolarity adjusted to 265 mOsm). For two-photon imaging, brains were then transferred fresh to 35mm imaging dishes and pinned to sylgard squares with tungsten wire. Ex vivo two photon imaging was performed on a Bruker Investigator using a 1.0 NA, 20x, water-dipping objective. Stacks were collected along the anterior-posterior axis with 1 micrometer spacing in Z and variable resolution in X/Y.

### Immunostaining and confocal microscopy

For immunostaining, after dissections brains were transferred to paraformaldehyde in PBS. Brains were fixed overnight at 4C in 1% PFA in PBS or for 25 minutes at room temperature in 4% PFA in PBS. In general, the 4% PFA method preserves cellular structures better, while the 1% PFA method allows better antibody penetration. After same-day (4%) or overnight (1%) fixation, brains were washed 3 times in PBS supplemented with 0.1% Triton-X-100 on a shaker at room temperature, blocked for at least 1 hour in PBS, 0.1% triton, 4% Normal Goat Serum, and then incubated for at least two overnights in primary antibody solution, diluted in PBS, 0.1% triton, 4% Normal Goat Serum. Primary antibody was washed 3x in PBS supplemented with 0.1% triton-x-100 on a shaker at room temperature, then brains were incubated in secondary antibodies for at least two overnights, diluted in PBS, 0.1% triton, 4% Normal Goat Serum. DAPI (1 microgram/mL) was included in secondary antibody mixes. Antibodies information can be found in the Key Resources Table Brains were mounted in 1x PBS, 90% glycerol supplemented with propyl gallate or SlowFade mounting medium in binder reinforcement stickers sandwiched between two coverslips. Samples were stored at 4C in the dark prior to imaging. The coverslip sandwiches were taped to slides, allowing us to perform confocal imaging on one side of the brain and then flip over the sandwich to allow a clear view of the other side of the brain. Scanning confocal stacks were collected along the anterior-posterior axis on a Leica SP8 with 1 micrometer spacing in Z and ∼150nm axial pixel size, using a 40x 1.3 NA oil immersion objective. Z dimension corresponds to anterior to posterior dimension of the central brain.

For Extended Data Fig. 8h and j, brains were stained and imaged as described in (Diamandi et al. 2024)^64^.

#### IHC with DPX mounting

In some cases, we sought to improve tissue clarity in IHC experiments using DPX mounting. To do this, brains were dissected in dissection saline, and then fixed in cold 4% PFA in PBS for 25min. Following fixation, brains were washed three times with 0.1% PBST (PBS containing 0.1% Triton X-100), then mounted onto a poly-L-lysine-coated coverslip. Blocking, primary antibody incubation and secondary antibody incubation were performed as described above. After secondary antibody incubation, the samples were washed with 0.1% PBST for three times, then subsequently post-fixed with 4% PFA for 2hrs at room temperature. After post-fixation, PFA was removed, and samples were washed four times with 0.1% PBST. The tissues were then dehydrated through a graded ethanol series (30%, 50%, 75%, 95%, and 2 × 100%), with each step lasting 5 minutes. This was followed by three washes in xylene (5min each). Finally, samples were mounted onto glass slides using DPX mounting medium. Mounted slides were air-dried for at least 2 days at room temperature before confocal imaging and stored at room temperature thereafter.

#### HCR RNA FISH

RNA fluorescent *in situ* hybridization (RNA-FISH) experiments were done as described by Ferreira et al., 2021 using Molecular Instruments HCR probe-sets, hairpins and reagents^142,143^. Brains were dissected in Schneider’s Drosophila Medium and fixed in cold 4% paraformaldehyde in PBS for 20 minutes at room temperature. Brains were rinsed 3x at room temperature in PBS supplemented with 1% triton-x-100 at room temperature then washed 3x with PBS supplemented with 1% triton-x-100 at room temperature. Samples were pre-hybridized in prewarmed Probe Hybridization Buffer (Molecular Instruments™) for 10 minutes at 37C and then incubated with HCR probes in Probe Hybridization Buffer overnight at 37C. The next day, they were washed with pre-heated Probe Wash Buffer (Molecular Instruments™) then washed 2 times in 5xSSCT (5X sodium chloride sodium citrate supplemented with 0.1% Tween-20). Samples were then preamplified in Amplification Buffer (Molecular Instruments™) for 10-30 minutes at room temperature and then incubated with snap-cooled HCR hairpins (95C for 90 seconds and then placed in the dark at room temperature for at least 30 minutes) in the dark overnight at room temperature. Before washing the brains, 300ul 5XSSCT was added to the samples to make the solution less viscous. The brains were then washed three times at room temperature first in 5XSSCT, then in Probe Wash Buffer (Molecular Instruments™) and then again in 5XSSCT. The brains were then rinsed one in Nuclease-Free PBS to remove any detergent and then were mounted in 1x PBS, 90% glycerol supplemented with propyl gallate and imaged as described above.

In some cases, RNA fluorescent *in situ* hybridization (RNA-FISH) experiments were done as described in Duckhorn et al., 2022^145^.

#### Combined RNA FISH and Immunohistochemistry

RNA fluorescent *in situ* hybridization (RNA-FISH) and Immunohistochemistry experiments were done as described by Schwarzkopf et al., 2021 using Molecular Instruments HCR probe-sets, hairpins and reagents^144^. The protein detection stage was first, with care to avoid introducing RNases into the solutions. Brains were dissected in external saline (108 mM NaCl, 5 mM KCl, 2 mM CaCl2, 8.2 mM MgCl2, 4 mM NaHCO3, 1 mM NaH2PO4, 5 mM trehalose, 10 mM sucrose, 5 mM HEPES pH7.5, osmolarity adjusted to 265 mOsm) and fixed in 4% paraformaldehyde in PBS for 25 minutes at room temperature. Brains were then washed 3x with PBS supplemented with 0.1% triton-x-100 at room temperature, blocked in Antibody Buffer (Molecular Instruments) at 4C for ∼4hrs, and then incubated overnight at 4C in primary antibody solution in Antibody Buffer (Molecular Instruments). Samples were then washed 4X in PBS supplemented with 0.1% triton-x-100 at room temperature and incubated in secondary antibody solution made in Antibody Buffer (Molecular Instruments). For some experiments we used the Molecular Instruments initiator labeled secondary antibody to amplify GFP signal (Donkey Anti-Rabbit-B4 for use with amplifier B4) and in other cases we used standard secondary antibodies (Goat Anti-Rabbit-488). Samples were incubated at room temperature for 3 hours then were washed 5X in PBS supplemented with 0.1% triton-x-100 then 1X in 5XSSCT both at room temperature. Samples were then preamplified in Amplification Buffer (Molecular Instruments™) for 10-30 minutes at room temperature and then incubated with snap-cooled HCR hairpins (95C for 90 seconds and then placed in the dark at room temperature for at least 30 minutes) in the dark overnight at room temperature. The brains were then washed in 5XSSCT at room temperature for (1) 2X5min (2) 2X30min and (3) 1X5min. The samples were then mounted as described above.

Alternatively, for R19C05*^Pdfr^* clonally labeled brains, we found that GFP was visible after RNA FISH even without immunostaining enhancement. Thus for some RNA FISH experiments on this genetic background, we conducted RNA FISH with care to keep the samples dark, and then imaged enduring native GFP fluorescence to visualize clones.

### Dye filling of mAL neurons

Dye filling of mAL neurons was performed as previously described^27,146^. Dye filling electrodes were pulled using a Brown/Flaming puller (Sutter Instruments) and were guided to the mAL axonal tract using *fru*-GFP. Voltage pulses (50-V pulses, 0.5 ms) were used to electroporate dye into the cells.

### Image analysis & statistical considerations

The sexes were separated across experimental conditions because mAL neurons are sexually dimorphic. To count cell populations, we used genetically encoded fluorescence or antibody staining as indicated. We counted labeled somata in every third slice in the stack (every third micron along the A-P axis), with reference to DAPI to distinguish individual cells from one another. Effect of mutations on axons and dendrites were done in a binary fashion due to the overt changes on brain structures induced by our manipulations, which also made it difficult for the researchers performing quantification to be blinded to experimental conditions. However, analysis was performed blind to the goals of the experiment when possible.

## Key resources tables

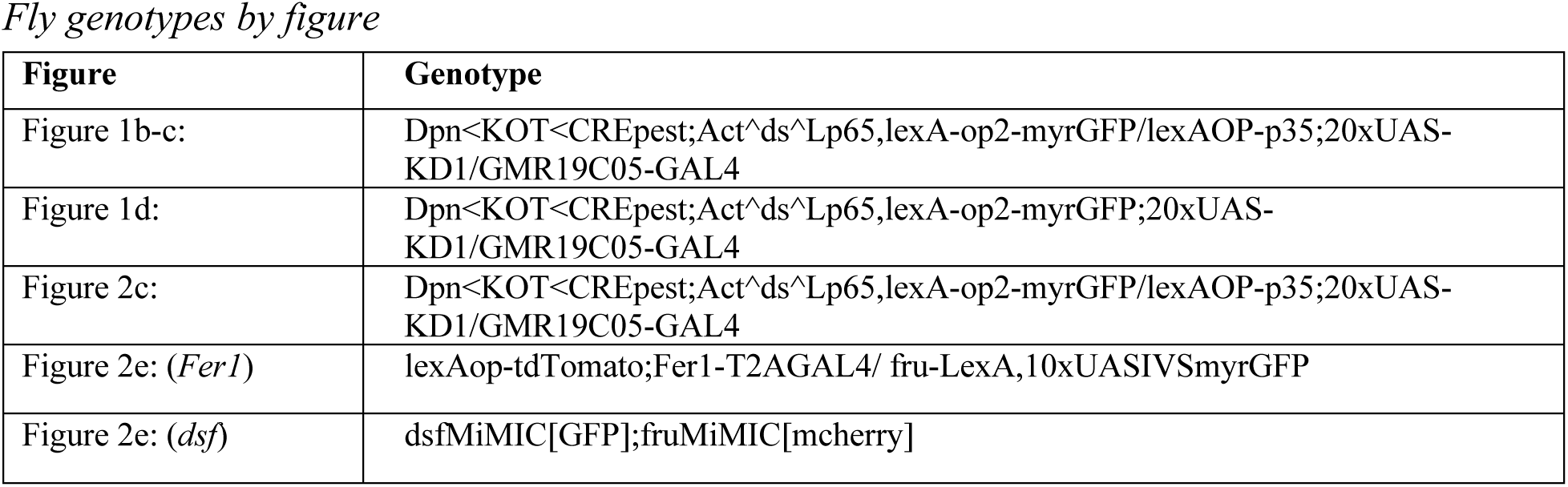
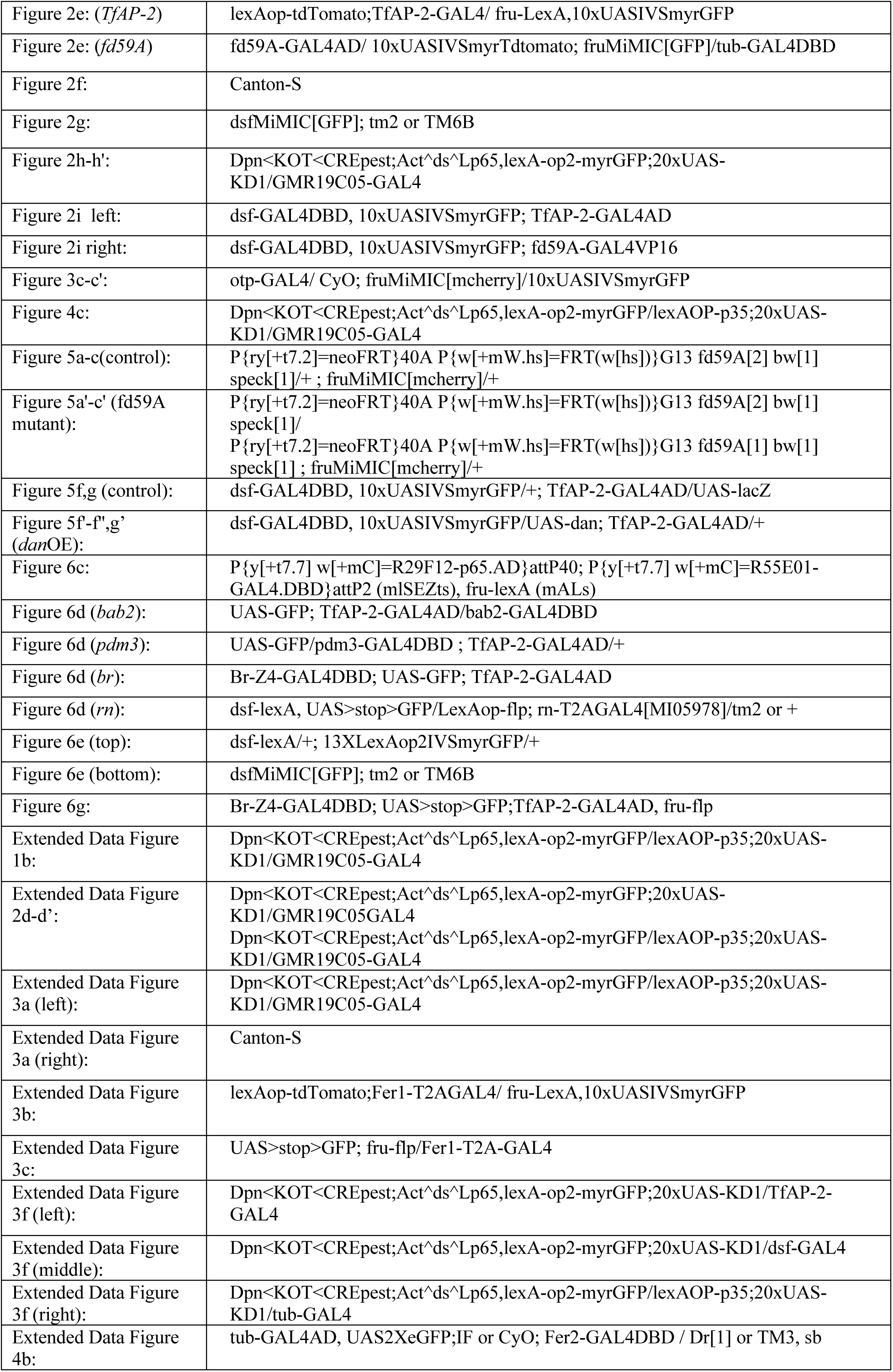
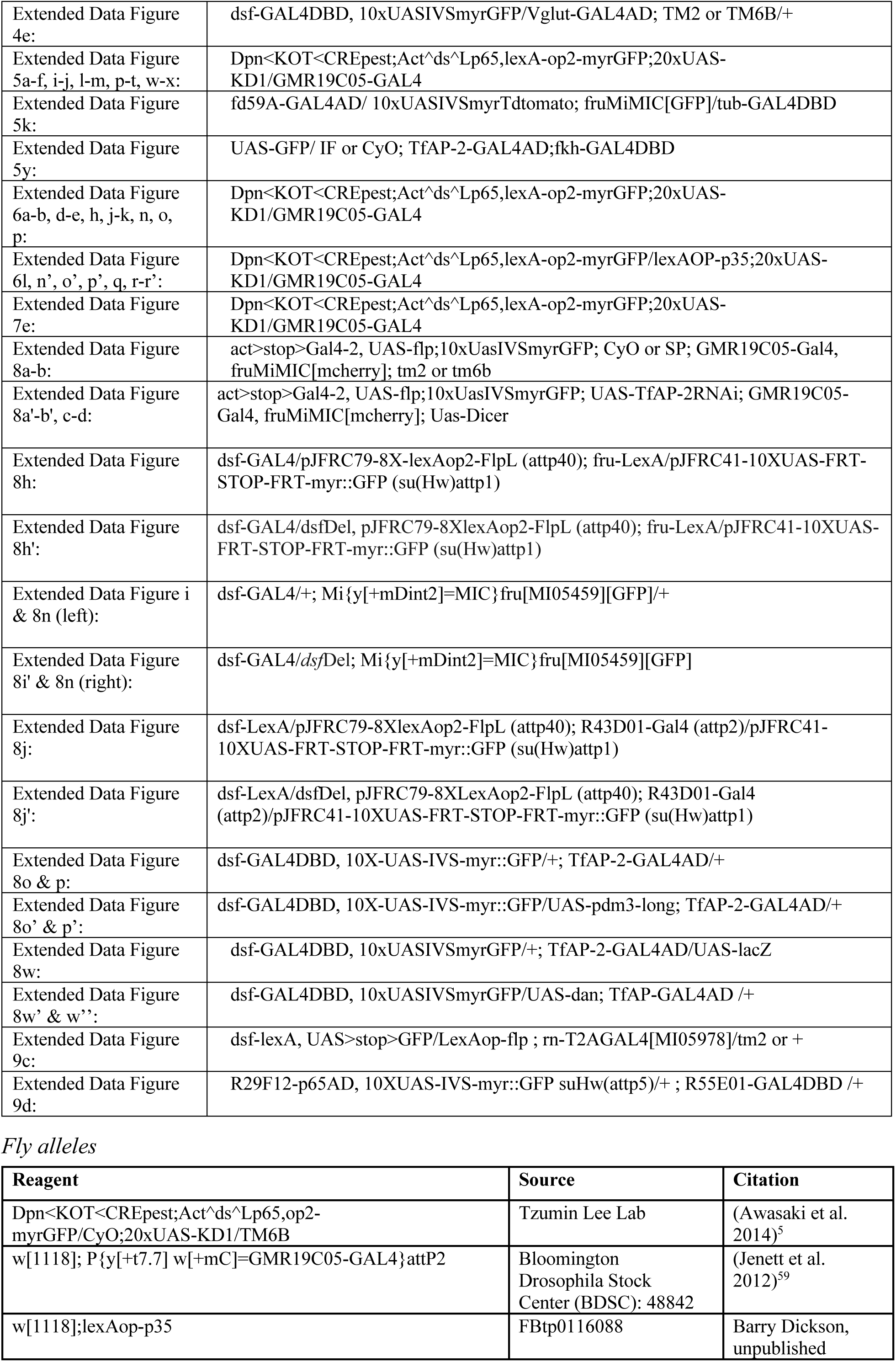
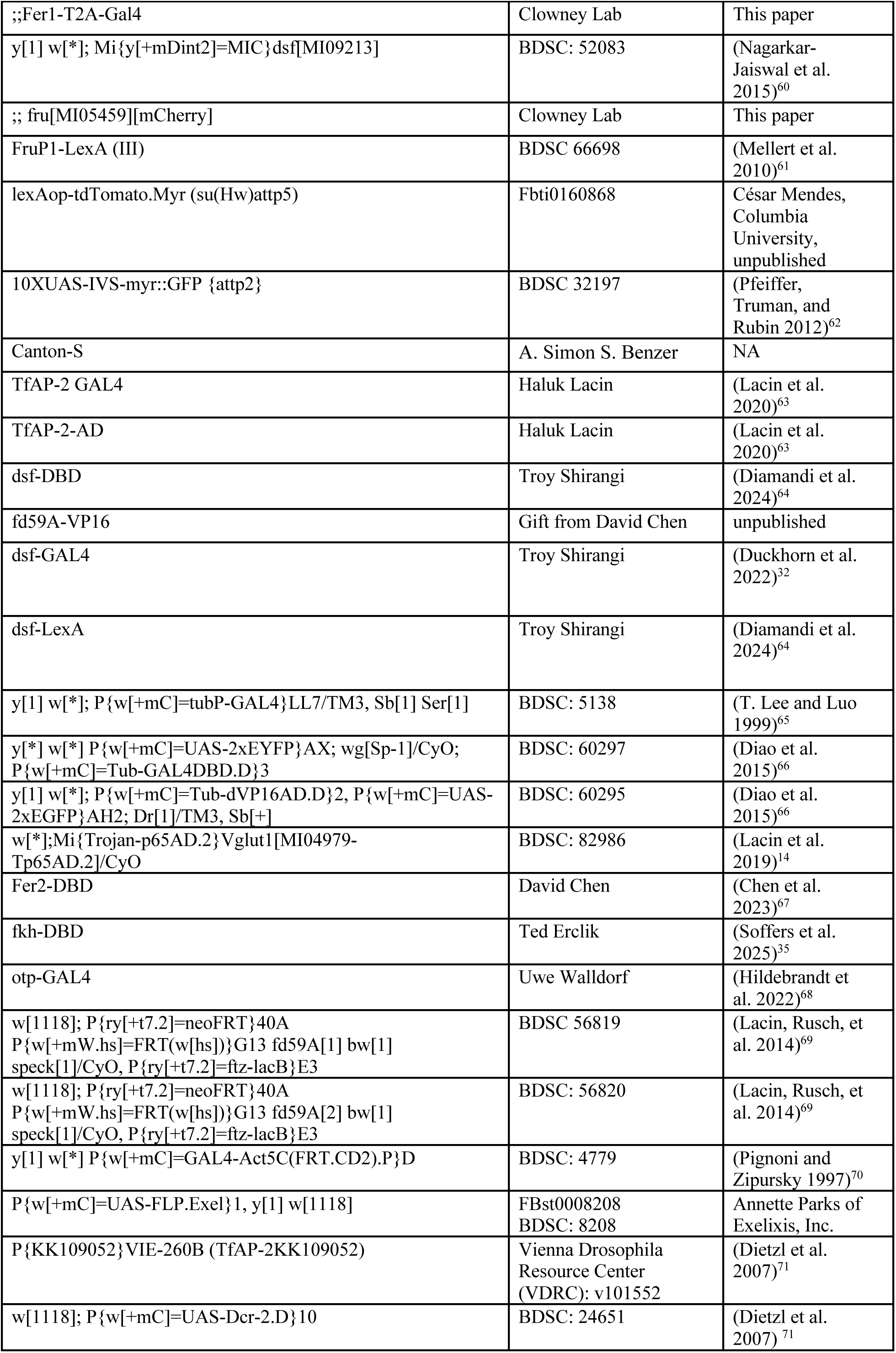
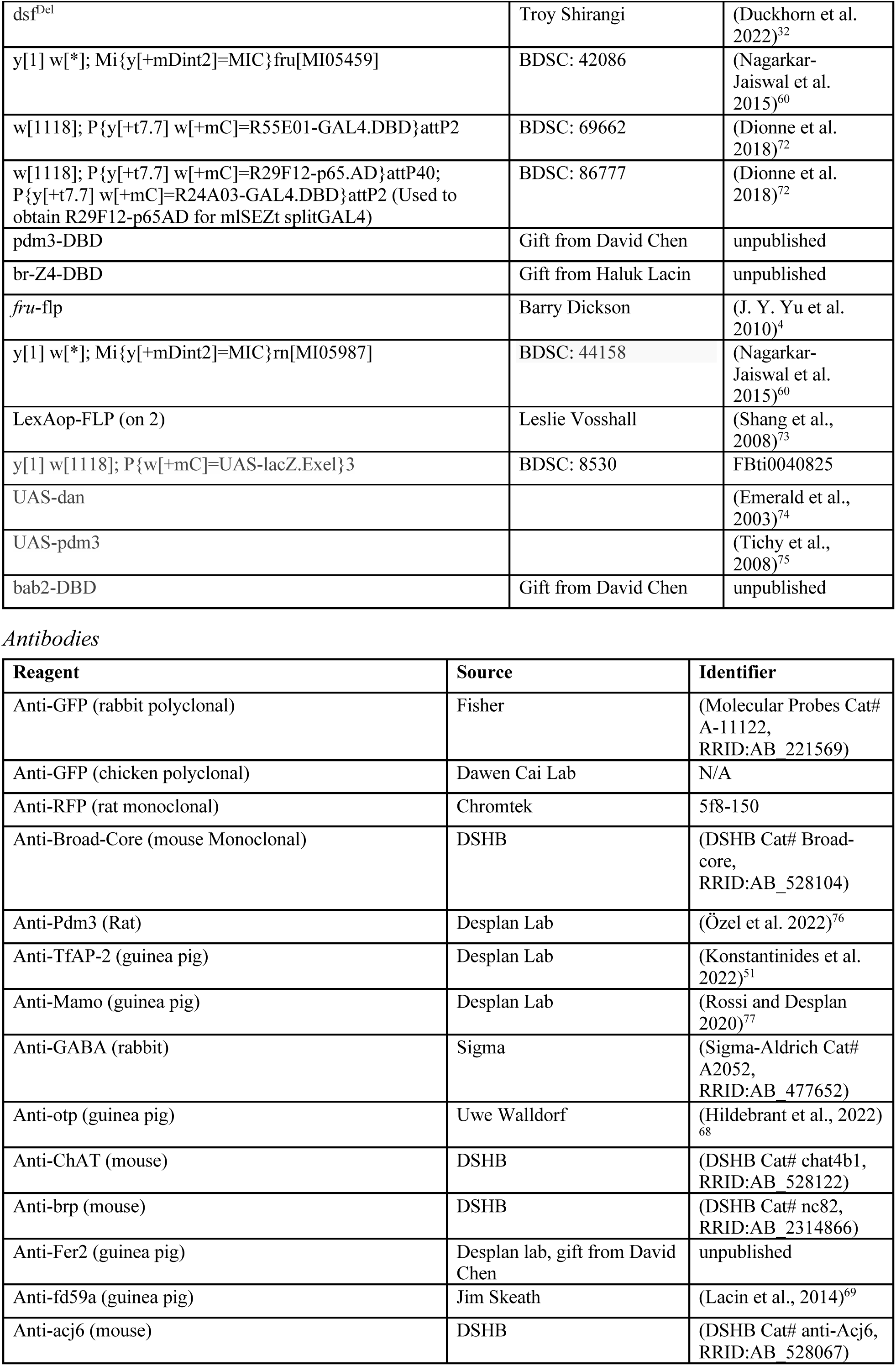
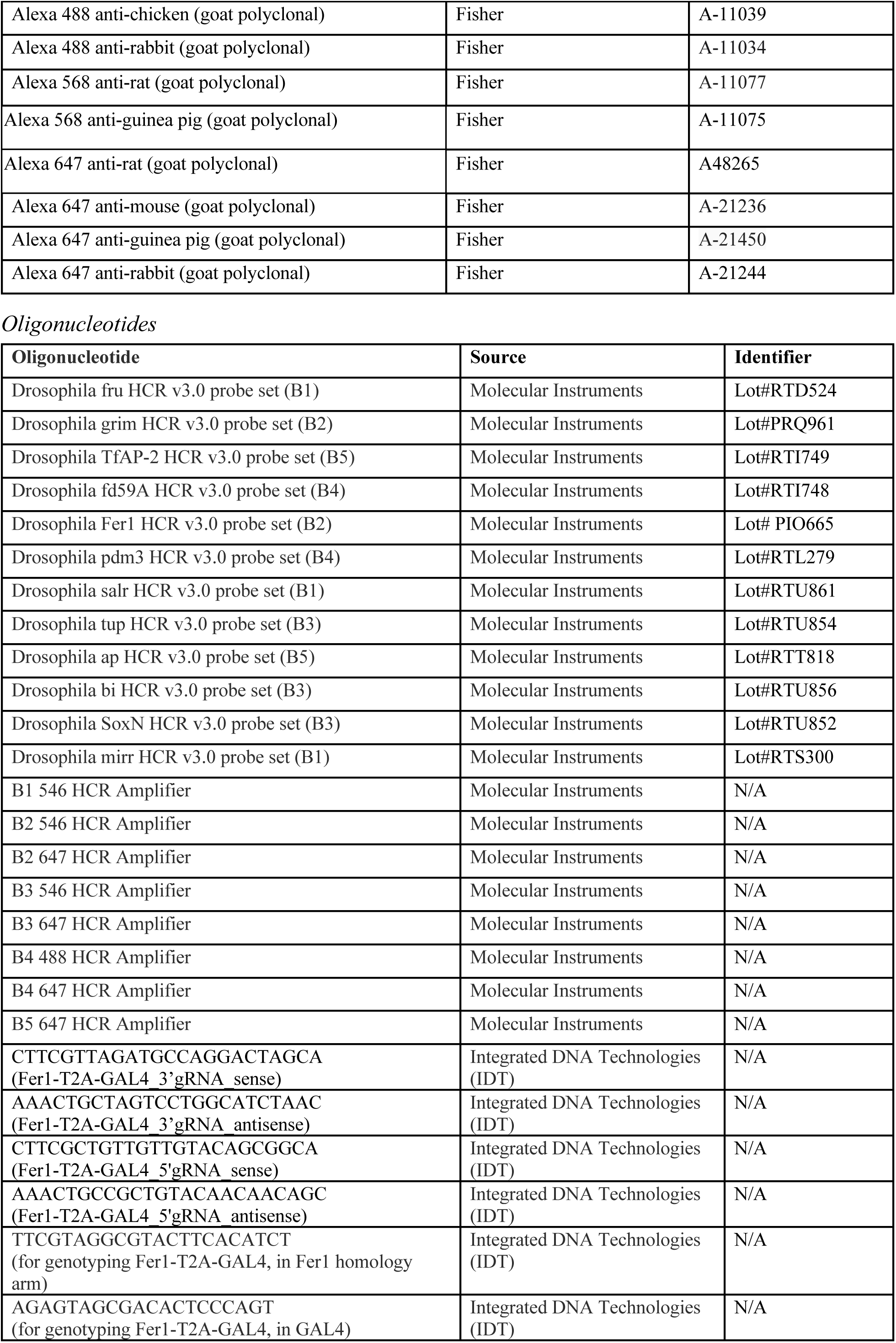
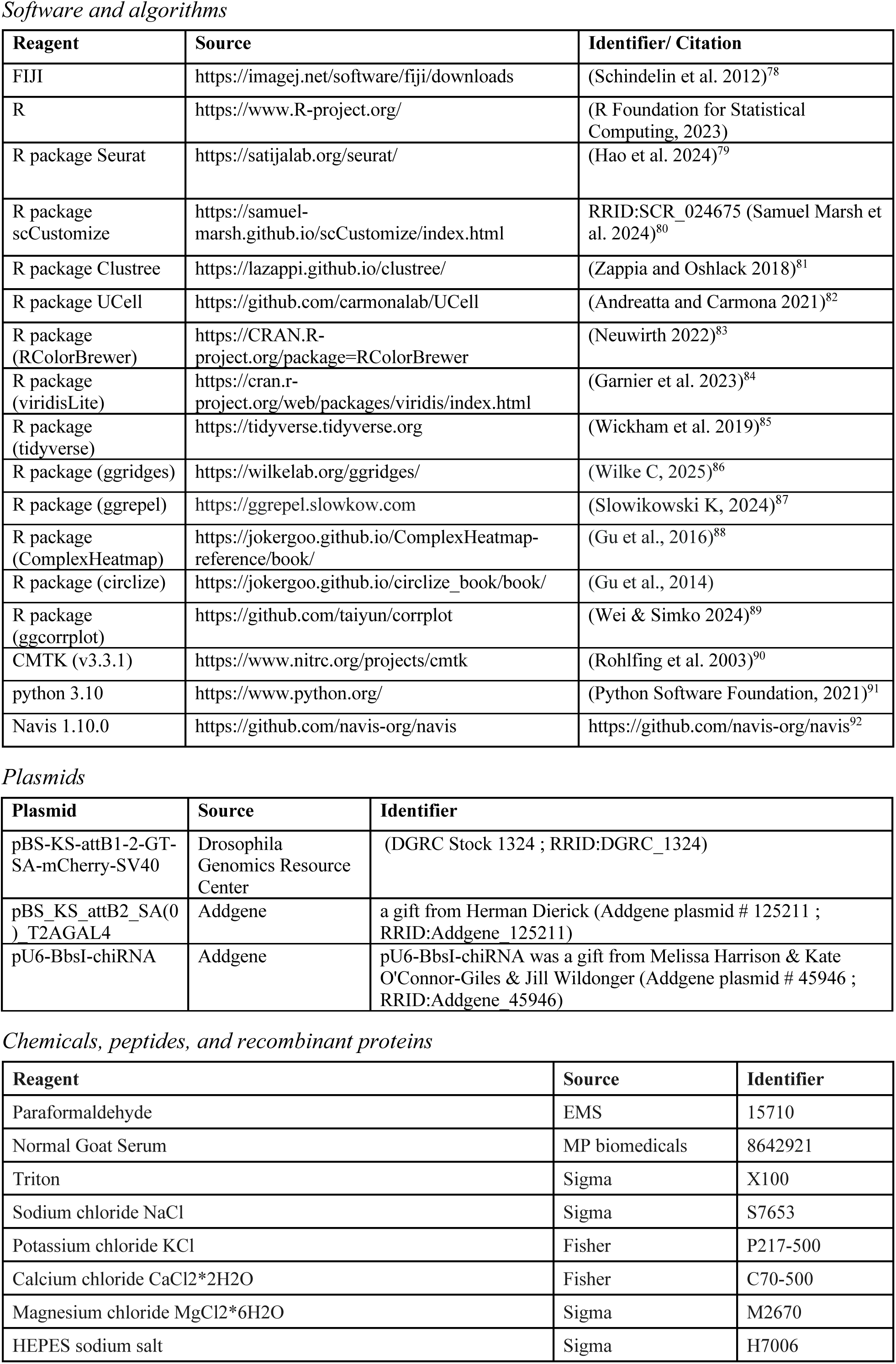
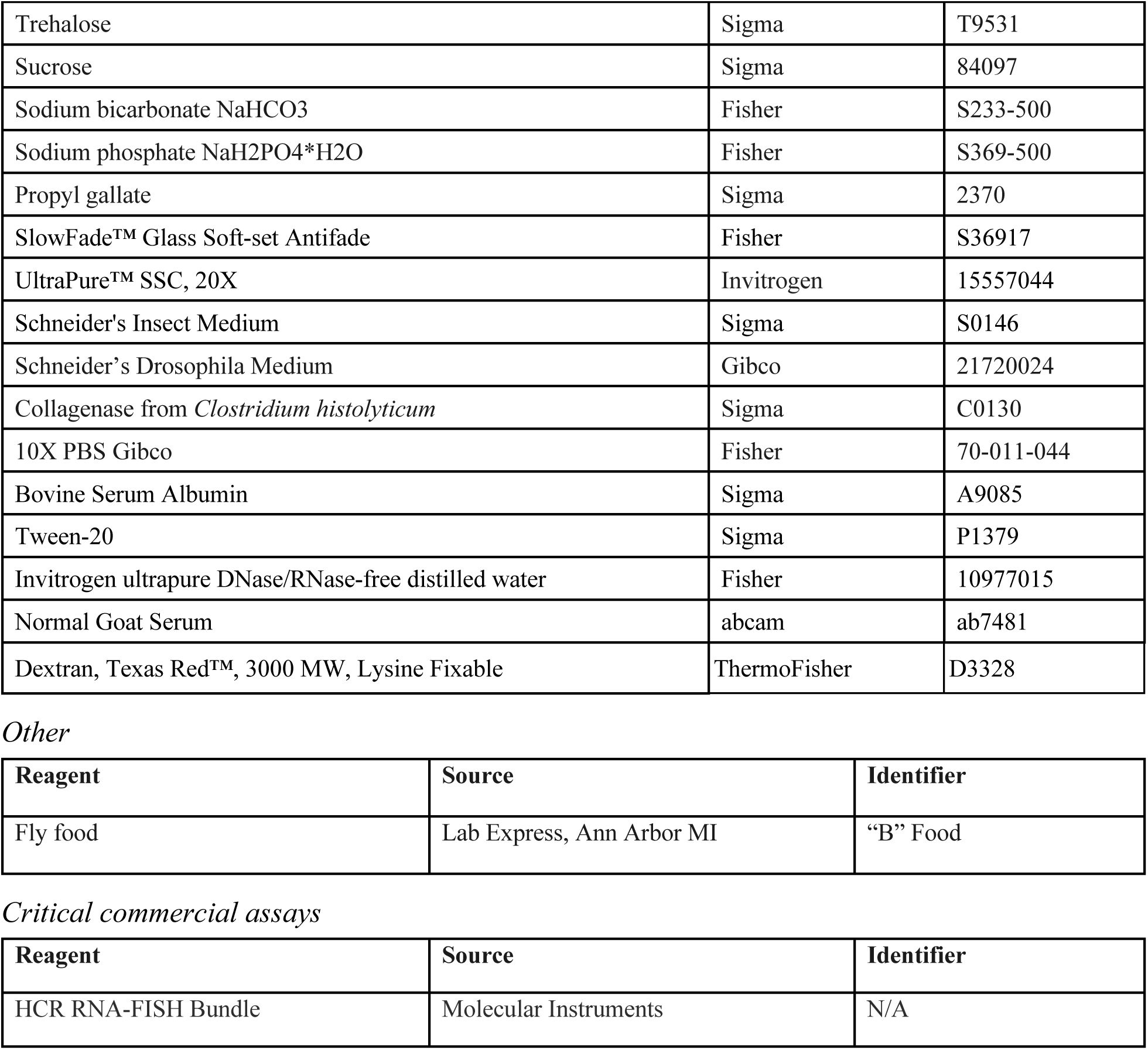

## Code Availability Statement

This paper did not generate custom code.

## Supplemental figures

**Extended Data Figure 1 (related to figure 1):**
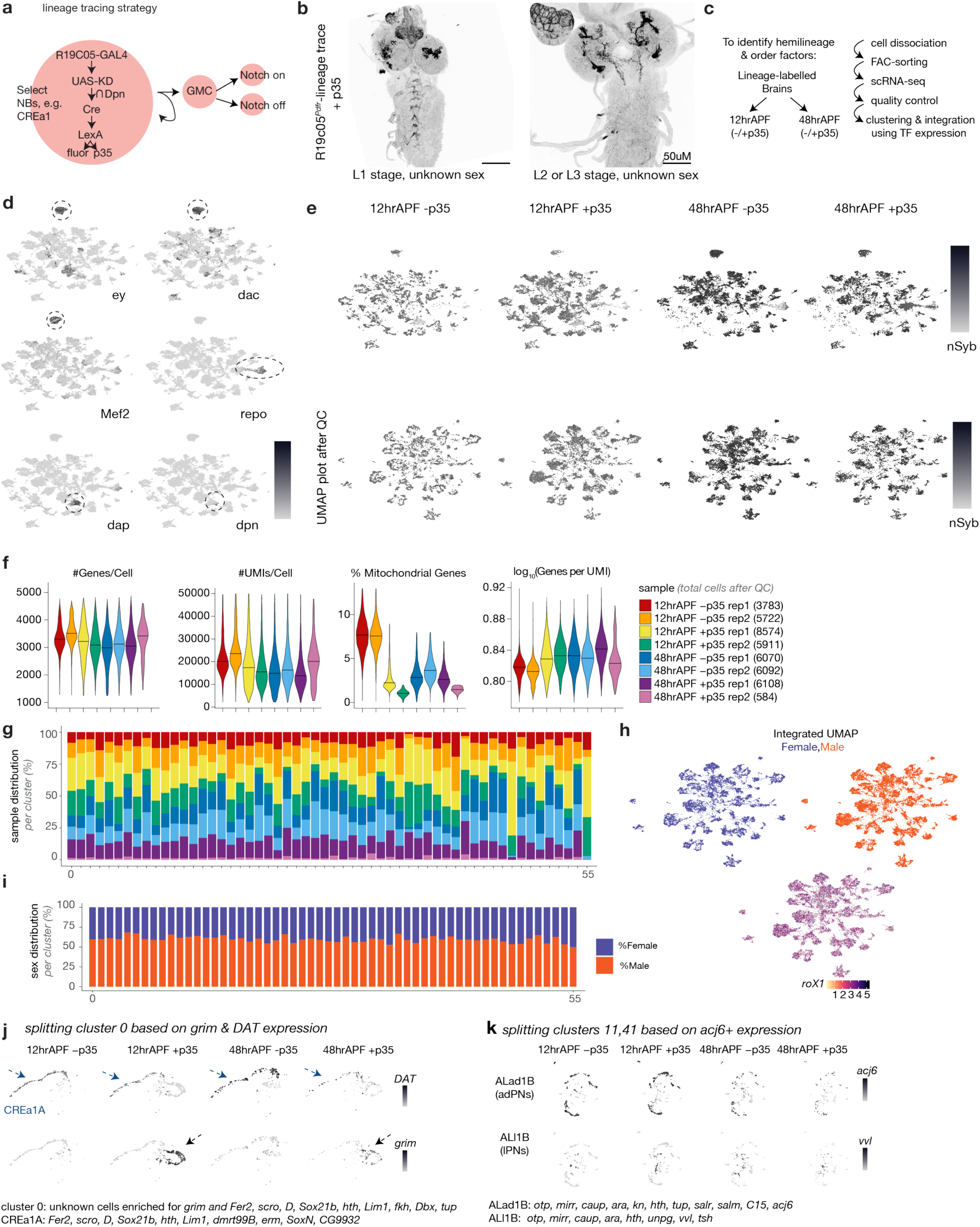
scRNAseq analysis of cerebral lineages. The R19C05 fragment of *pdfr* drives expression of GAL4 in CREa1 and other neuroblasts^22^. GAL4 drives KD recombinase. The intersection of KD and the neuroblast-specific *deadpan* promoter activates Cre. Cre excises a stop codon upstream of LexA. This results in permanent LexA expression in the NB and its daughters, which can be used to drive expression of effectors, including fluorophores or the viral caspase inhibitor p35. Genetic strategy adapted from Awasaki et al. 2014^5^. **b**, R19C05 Clonal labeling strategy in early larval stages (L1) vs L2/L3 determined by mouth hooks. **c**, scRNAseq and analysis workflow. **d**, UMAP feature plots of pre-filtered scRNA-seq data from R19C05-labeled lineages. *Ey*, *dac* and *Mef2* mark Kenyon cells, *repo* marks glia and *dpn* and *dap* mark progenitor cells. **e**, UMAP feature plots of nSyb expression split across timepoints (12hrAPF and 48hrAPF) and conditions (+/-p35) before filtering *nSyb*-low cells (top) and after (bottom). **f**, The number of genes, UMIs, percent mitochondrial gene expression and the “library complexity” (log10 genes/log10 UMI) across cells for each library after filtering. The final number of cells from each dataset is between parentheses. **g**, The proportion of cells coming from each of our 8 datasets for every cluster. **h**, Sexing cells using *lncRNA:roX1* and *lnRNA:roX2* and contribution of male and females cells across the integrated UMAP. **i**, The proportion of cells coming from males versus females based on the expression of *lncRNA:roX1* and *lnRNA:roX2*. **j**,**k**, Basis and method for splitting clusters 0 (**j**) and 11 and 41 (**k**) into distinct hemilineage clusters; rationale is described in the Methods and in Supplementary Table 1.

**Extended Data Figure 2 (related to figure 1):**
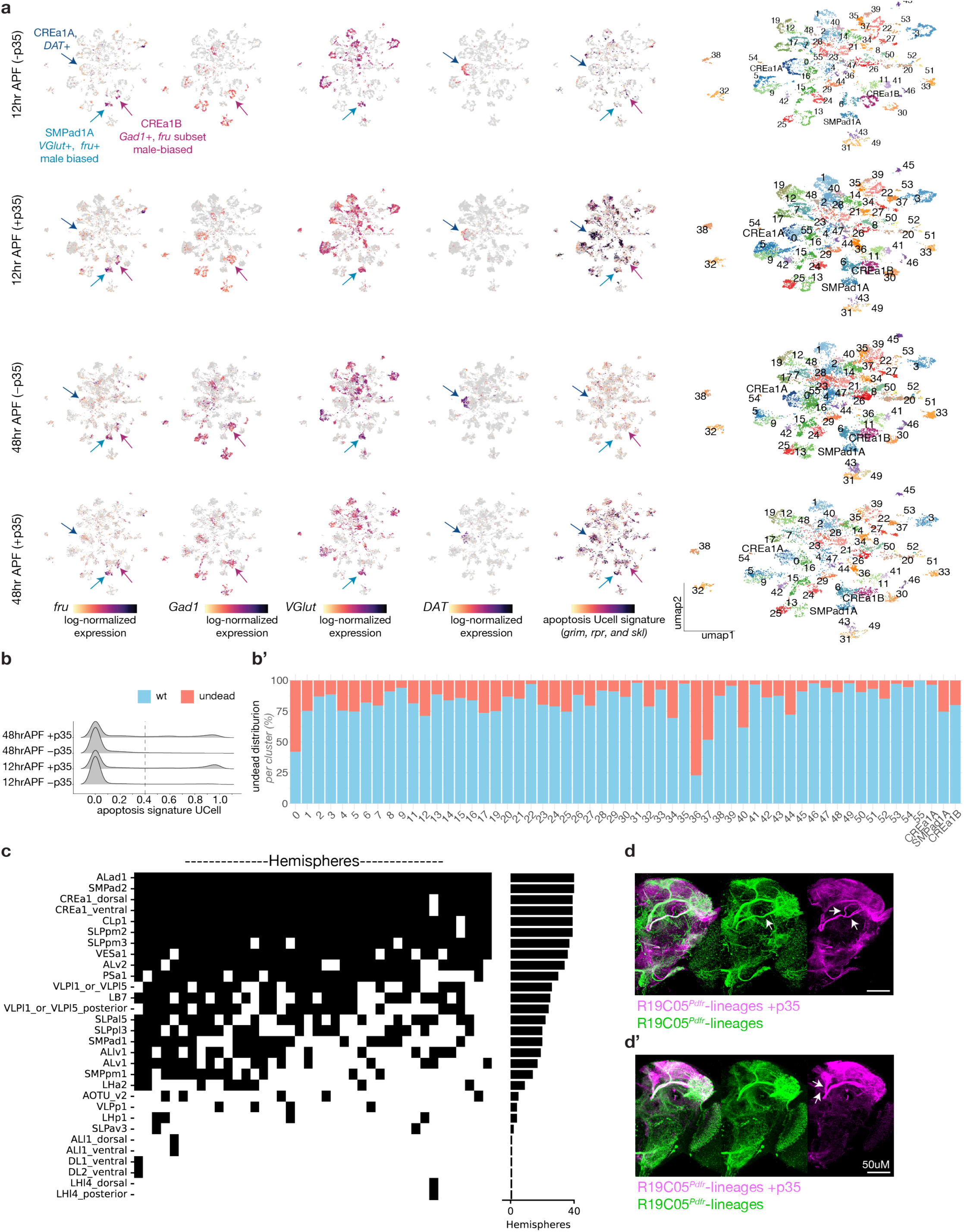
Characterization of transcriptional clusters and anatomic lineages. **a**, UMAP feature plots of scRNA-seq data from R19C05-labeled lineages, split across timepoints (12hrAPF and 48hrAPF) and conditions (+/-p35). CREa1A, B and SMPad1A marked by arrows. At right, contribution of each library type to each cluster. **b**, Distribution of UCell gene enrichment scores for apoptotic genes *grim*, *rpr*, and *skl* across cells from libraries of each type. Cells with an apoptosis UCell score above 0.4 only arise from libraries in which we blocked the completion of apoptosis (+p35); we thus designate cells above this threshold as “resurrected.” **b’**, Proportion of resurrected cells in each cluster from the UMAP. Three clusters, 0, 36, and 37, are dominated by resurrected cells. **c**, Census of lineage/hemilineages labeled by R19C05*^Pdfr^* lineage trace. We registered 10 male and 10 female labeled brains to the JFRC18 common brain template and annotated which tracts were labeled in each hemisphere (columns) to identify hemilineages (rows). Pairs of hemilineages that arise from the same neuroblast are always labeled together. At right, sum of how often each hemilineage was observed in the census. **d**, **d’**, Identification of ectopic, resurrected tracts that are only observed in the R19C05*^Pdfr^* lineage traced clones in the presence of p35. Example hemispheres with and without p35 were registered to the common brain template and overlaid. Arrows in (**d**) highlight the addition of ectopic tracts similar to that naturally produced by ALv1, and in (**d’**) addition of a medially-extending tract from the SMPpm1 lineage. Note that additional differences between the individual hemispheres shown are due to inconsistent clonal induction by R19C05*^Pdfr^* lineage trace.

**Extended Data Figure 3 (related to figure 2):**
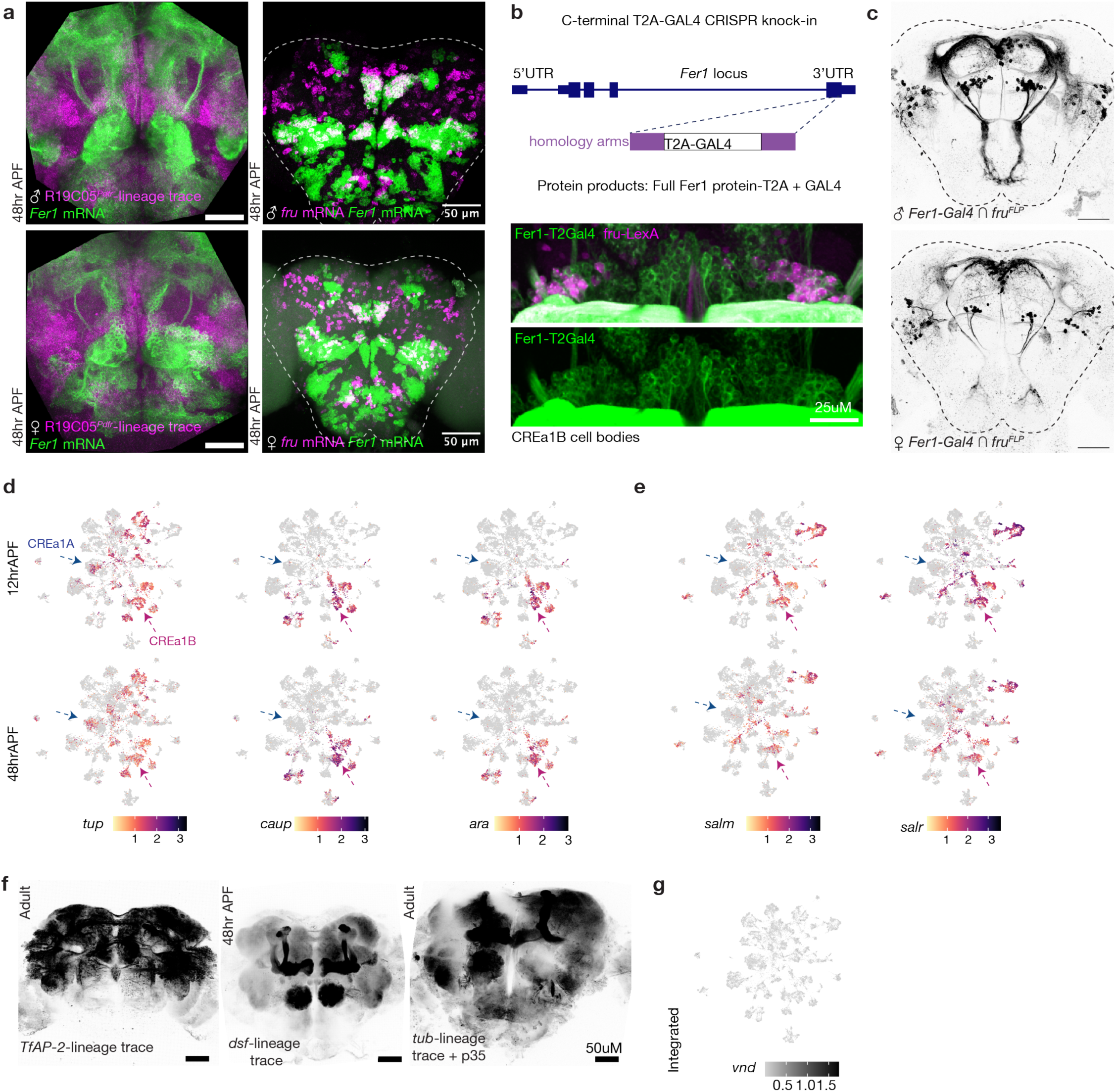
Analysis of additional transcription factors in CREa1B hemilineage or CREa1 neuroblast. **a**, (Left) R19C05*^Pdfr^*clonal labeling combined with RNA FISH for *Fer1* at 48hrAPF. (Right) RNA FISH for *Fer1* and *fru* (close-up view of somata shown in Figure 2). **b**, (Top) CRISPR knock-in strategy to C-terminally tag Fer1 with T2A-GAL4. (Bottom) CREa1B cell bodies labeled by the Fer1-GAL4 in green and *fruitless*-expression labeling in magenta with *fru*-lexA. **c**, Two photon image of neurons labeled by genetic intersection of Fer1-T2A-GAL4 with *fru*-Flp. **d**, **e** UMAP plots of additional TFs expressed in CREa1B, displaying the log-normalized expression in cells split by time (12 and 48hrAPF). **f**, Lineage labeling using the Awasaki strategy driven by dsf-GAL4 (left), TfAP-2-GAL4 (middle) and tub-GAL4 with the apoptotic block (lexAop-p35) (right). Fer1-T2A-GAL4 and TfAP-2-intersect-dsf-split-GAL4 did not produce lineage labeling. **g**, UMAP plot displaying the log-normalized expression of *vnd*, showing that *vnd* is not expressed in postmitotic neurons.

**Extended Data Figure 4 (related to figure 2):**
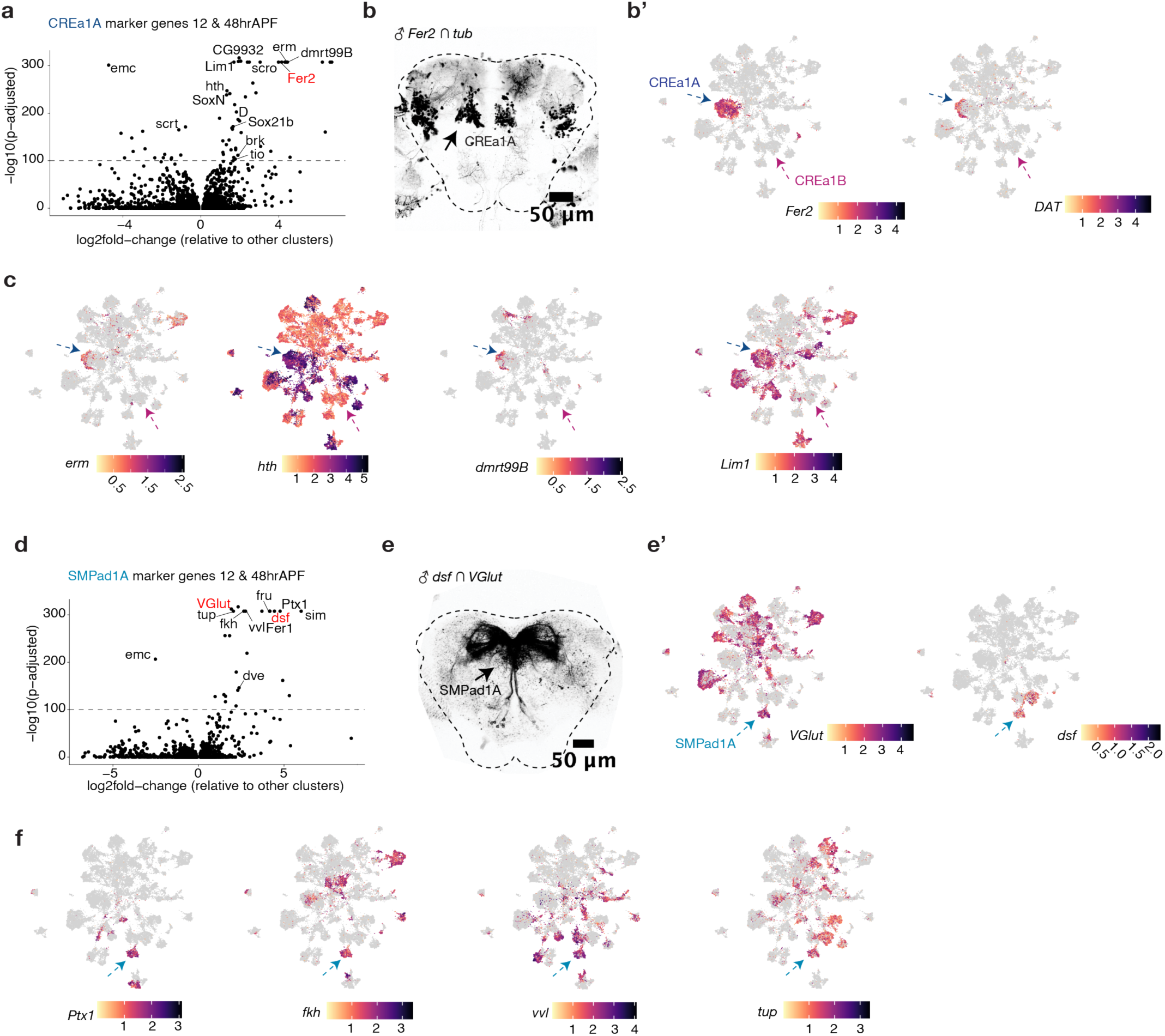
Matching the CREa1A and SMPad1A hemilineages to transcriptional clusters. **a**. TFs enriched in the putative CREa1A cluster. **b**, **b’** The only hemilineage within the R19C05*^Pdfr^* clone set that expresses *Fer2* is CREa1A. (*tubulin* splitGAL4 is universally expressed and used to convert another splitGAL4 allele into a “regular” GAL4.) **c**, UMAP Featureplots integrated across time showing hemilineage factors in CREa1A. **d**. TFs enriched in the putative SMPad1A cluster. **e**, **e’** The only glutamatergic hemilineage within the R19C05*^Pdfr^* clone set that expresses *dsf* is SMPad1A. **f**, UMAP Featureplots integrated across time showing hemilineage factors in SMPad1A.

**Extended Data Figure 5 (related to figure 3):**
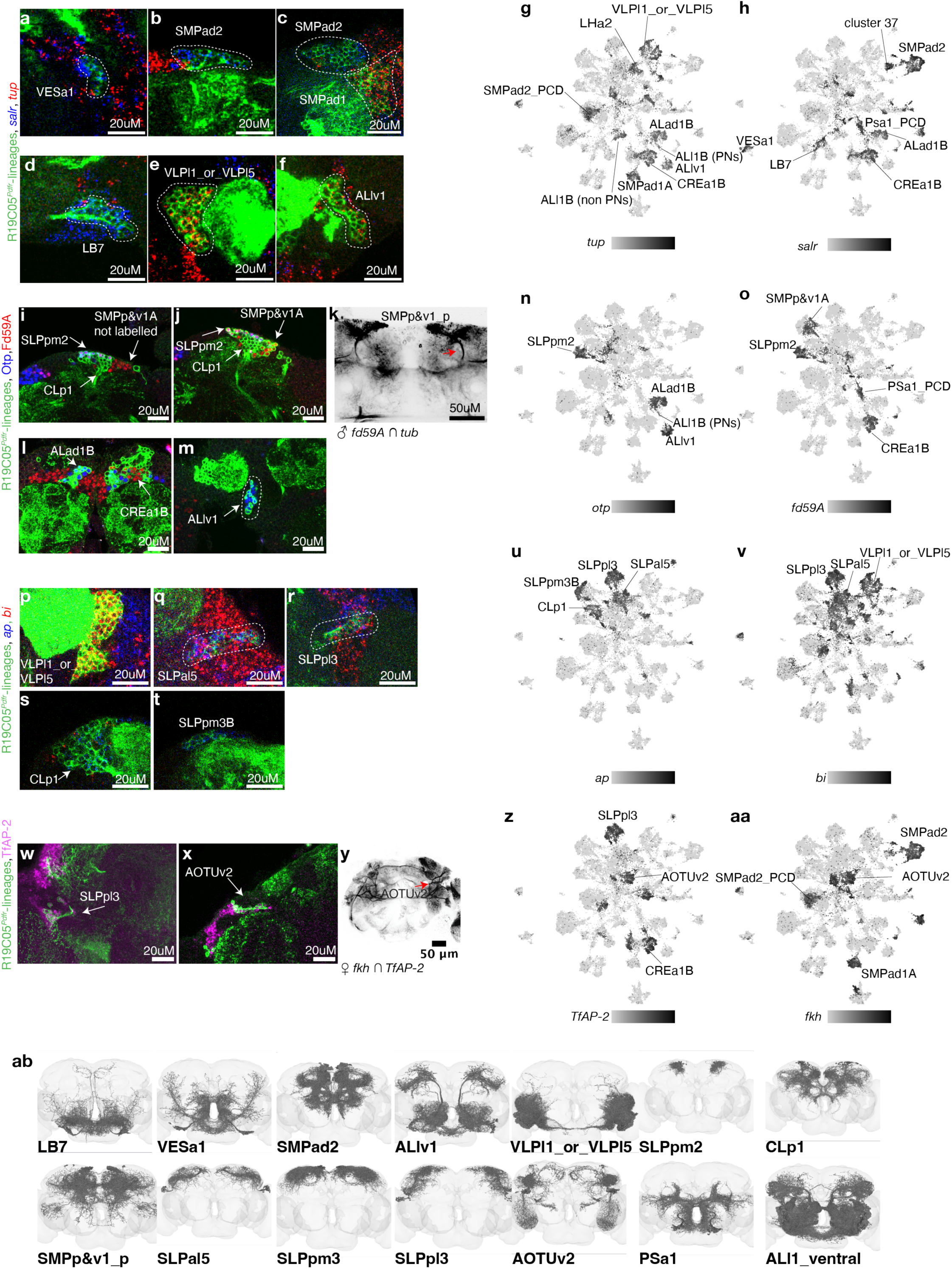
Using histology to match anatomic hemilineages to scRNAseq clusters, 1. **a-aa**, In each section of panels, immunostaining or HCR RNA FISH is performed on top of the R19C05*^Pdfr^* clone set, as indicated, allowing identification of anatomic hemilineages that express the TFs of interest. At right, UMAP plots showing those same TFs and positively identified hemilineages that express them. **ab**, anatomy of matched hemilineages as annotated in FAFB^15,96^. Additional description is provided in the Methods, in Supplementary Tables 1 and 3, and in Extended Data Figure 6.

**Extended Data Figure 6 (results related to figure 3):**
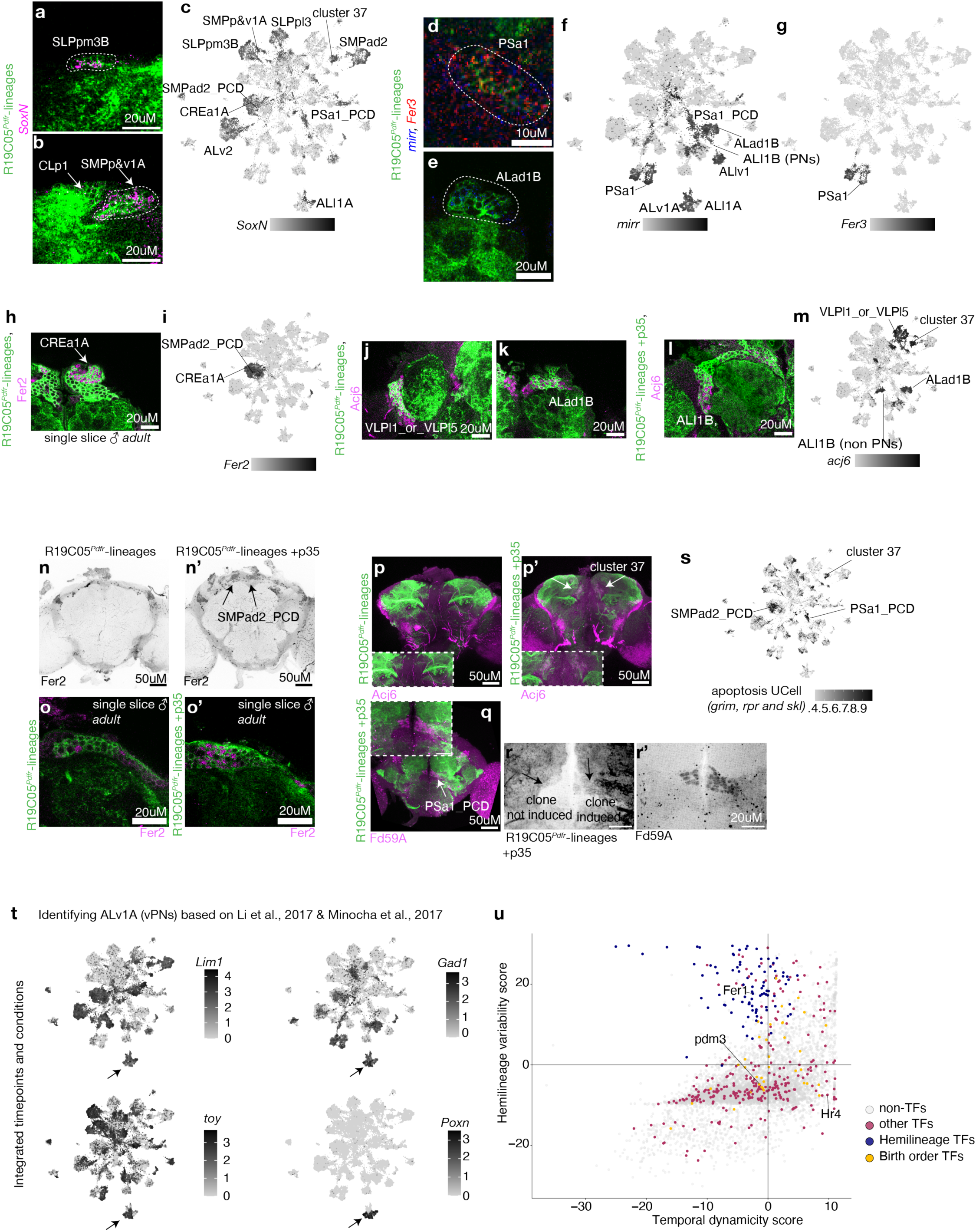
Using histology to match anatomic hemilineages to scRNAseq clusters, 2. **a-m**, In each section of panels, immunostaining or HCR RNA FISH is performed on top of the R19C05*^Pdfr^*clone set, as indicated, allowing identification of anatomic hemilineages that express the TFs of interest. UMAP plots show those same TFs and positively identified hemilineages that express them. **n**, **o**, Identification of a resurrected hemilineage that is sister to SMPad2. SMPad2 is labeled in 100% of our R19C05*^Pdfr^*lineage traced brains. In the natural genotype, scant Fer2-positive cells are observed in this region of the brain, while many cells are observed in the p35 genotype. The ectopic transcriptional cluster expressing *Fer2* is shown in (**i**). This cluster corresponds to cluster 0 in Extended Data Figure 2b’, which is enriched for resurrected cells. **p**, **p’** Identification of resurrected cluster 37; Acj6 only appears in this part of the brain in the +p35 genotype. While this anatomic set corresponds to cluster 37 in the UMAP, we have not determined which neuroblast it arises from or its sister hemilineage. **q**, **r**, **r’**, Identification of a resurrected sister hemilineage to PSa1. The brain shown is of the p35 genotype, but the PSa1 lineage clone is induced only in the right hemisphere. Absent PSa1 clone induction, or in non-p35 brains, there are just a few Fd59a cells in this regions, which correspond to primary neurons sister to PSa1, called aDT7. When p35 is expressed, many more Fd59a cells are labeled. The PSa1_PCD cluster is #36 in Extended Data Figure 2b’, which is enriched for resurrected cells. Expression of *fd59a* in this cluster can be seen in Extended Data Figure 5l. **s**, Apoptosis UCell highlights these three resurrected hemilineages. Additional description is also provided in the Methods and in Supplementary Tables 1 and 3. **t**, Use of *Lim1*, *toy*, and *PoxN* allow identification of ALv1A. **u**, Statistical analysis of the variation in expression of hemilineage, birth order, and other TFs across developmental time (x axis, as described by Jain et al., 2022) or across clusters (y axis). Examples of a hemilineage TF (Fer1), a birth order TF (pdm3) and a maturation TF from Jain et al (Hr4) are highlighted. We defined hemilineage TFs as having high variability across clusters but stable expression over developmental time. Birth order TFs are variable within, but not across hemilineage clusters. To qualify, we required that they be expressed in subsets of cells within hemilineage clusters at both our developmental timepoints, but their expression levels can vary. TFs outside these TFs follow the same distribution as the rest of the genes in the genome.

**Extended Data Figure 7 (results related to figure 4):**
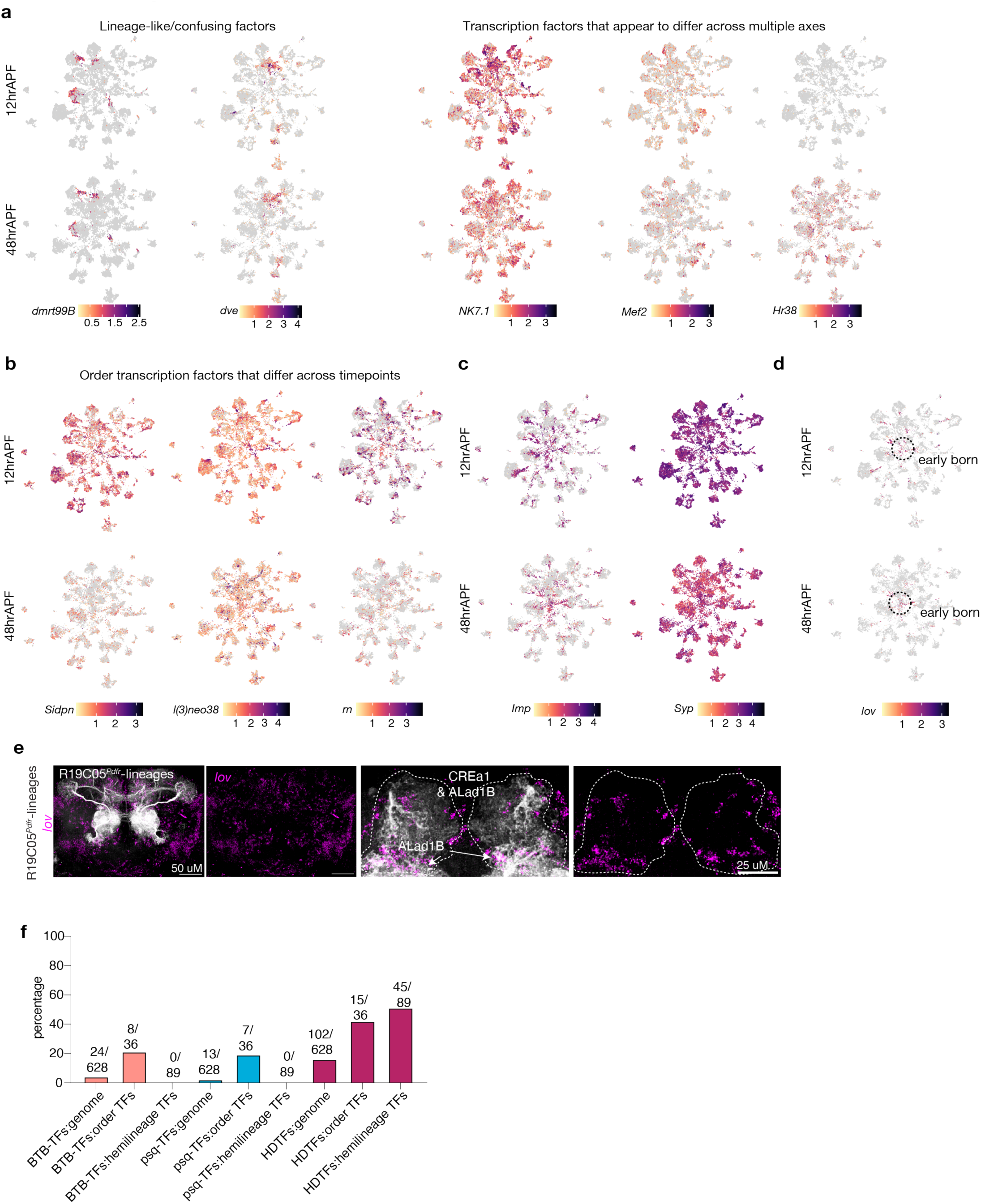
Additional description of putative birth order and other TFs. **a**, Transcription factors with “weird” expression, i.e. varying along more than one developmental/diversification axis. **b**, While we required that putative birth order TFs be expressed in subsets of cells within hemilineage clusters at both our developmental timepoints, expression levels could vary across timepoints, as in the examples shown here. **c**, Expression of temporally associated RNA binding proteins *Imp* and *Syp*. In cerebral Type I lineages, *Imp* transcription resembles that of a birth order factor, while *Syp* transcription is pervasive. As these proteins regulate each other’s translation, Syp protein expression is likely to be more specific. **d**, Identification of a cluster of cells enriched for *lov*. This cluster is enriched for *Imp* and likely to contain mixed, early-born neurons (described in Methods). **e**, Assesment of the expression pattern of *lov* within clones. Consistent with a role as a birth order factor, *lov*-expressing cells are a small set of each clone. **f**, Distribution of BTB, pipsqueak, and homeo-domains among birth order and hemilineage TFs compared to the genome.

**Extended Data Figure 8 (related to figure 5):**
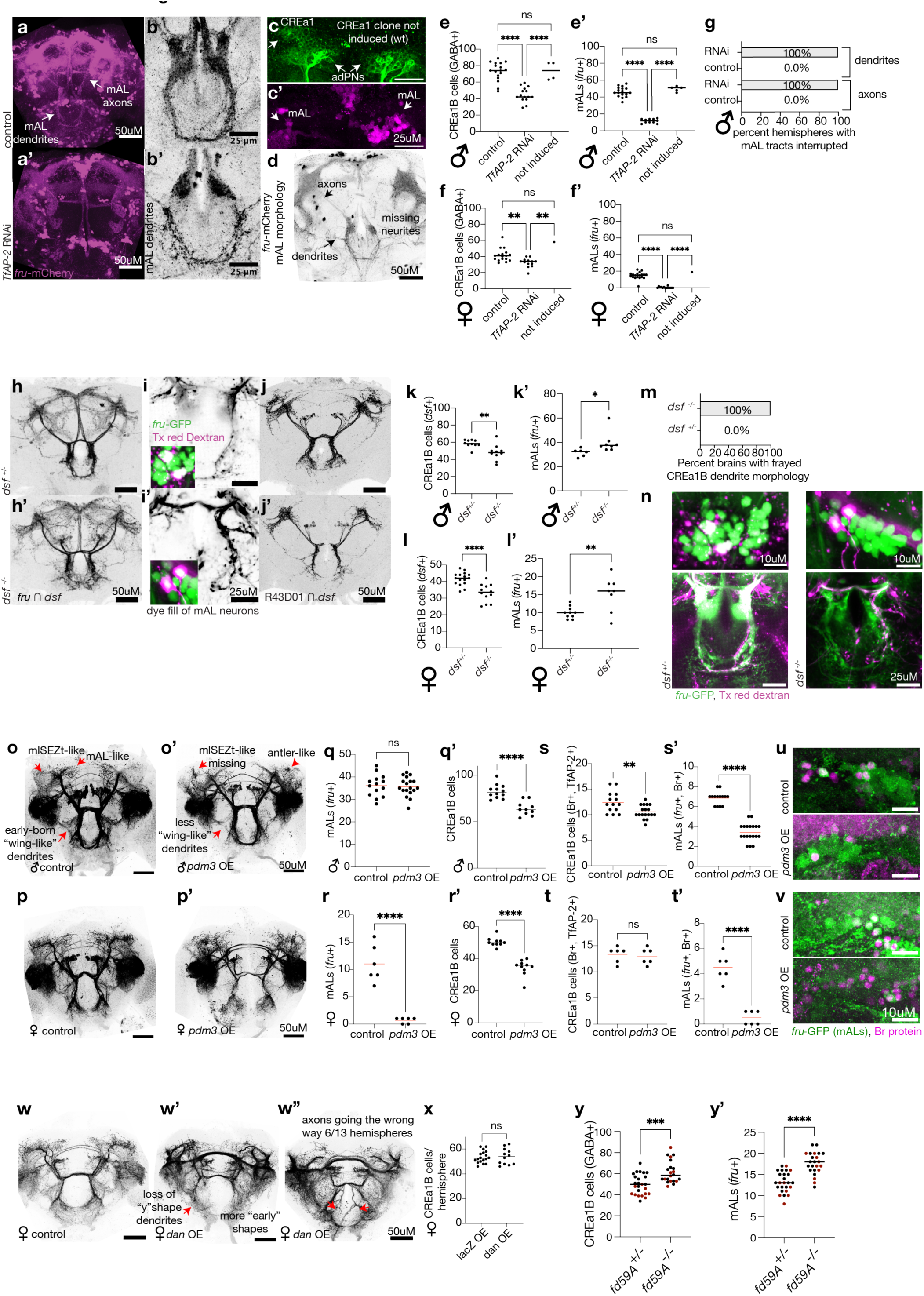
Phenotypes associated with alteration of additional hemilineage or birth order TFs. **a-c**, Morphology of mAL neurons labeled by *fru*-mCherry in controls versus *TfAP-2* RNAi clones. (**b**) Displays mAL neuron dendrites zoomed in, while **c** and **d** show the soma (**c**) and axons (**d**) of control and *TfAP-2* RNAi clones within the same brain. The CREa1B soma are labeled by R19C05-lineage labeling and mAL neurons soma and morphology are labeled using *fru*-mCherry. Arrows, mAL axons or dendrites. **e, e’, f, f’**. The number of total CREa1B neurons (GABA+). or mALs per hemisphere (*fruitless*-expressing). Significance, one-way ANOVA. “Non-induced” refers to hemispheres from animals of the *TfAP-2* RNAi genotype in which clonal labeling of CREa1 was not induced, while “control” refers to the genotype lacking the RNAi allele. **g**, The qualitative effect of the loss of *TfAP-2* on mAL axonal and dendritic morphology. **h**, Morphology of mAL neurons labeled by intersection of *dsf* and *fru* in controls versus *dsf* mutants. **i**, Texas red Dextran electroporated into the mAL axonal tract in control and *dsf* mutants; insets showing somata indicate that dye fill is restricted to *fru*-expressing cells in the lineage. Representative of three or more experiments per genotype. **j**, The intersection of R43D01 and *dsf* labeling in control and *dsf*-mutant brains. This intersection includes *fruitless*-expressing and non-expressing CREa1B neurons. **k, k’, l, l’**. The number of total CREa1B neurons (*dsf*+) or mALs per hemisphere (*fruitless*-expressing). Significance: unpaired t-test. **m**, Summary of the qualitative effect of losing *dsf* on dendritic morphology. **n**, Additional images of the samples shown in (**i**). **o**, **o’**, **p**, **p’**, The SplitGAL4 intersections of *dsf* and *TfAP-2* in control versus Pdm3 overexpression in males and females. Red arrows highlight structures of CREa1B axonal and dendritic subtypes. q, q’, r, r’, : Number of total mAL neurons (*fruitless*-expressing) or total CREa1B cells (labeled by *dsf* intersect *TfAP-2* split-GAL4) per hemisphere in control of pdm3-overexpressing males and females. Significance: unpaired t-test. **s, s’, t, t’** Number of Br-expressing CREa1B neurons (based on TfAP-2 protein expression) per hemisphere and number of Br-expressing mAL neurons (*fruitless*-expressing) in male and female control and *pdm3* overexpression animals. **u, v** mAL cell bodies labeled with the *fru* transcriptional reporter (*fru*-GFP) in green and Br protein in magenta in male and female control and *pdm3* overexpression animals. **w, x** Images and quantification of female brains with Dan overexpressed under control of the SplitGAL4 intersection of *dsf* and *TfAP-2*. **y,y’**Quantification of CREa1B and mAL neurons in females control or mutant for *fd59a*.

**Extended Data Figure 9 (results related to figure 6):**
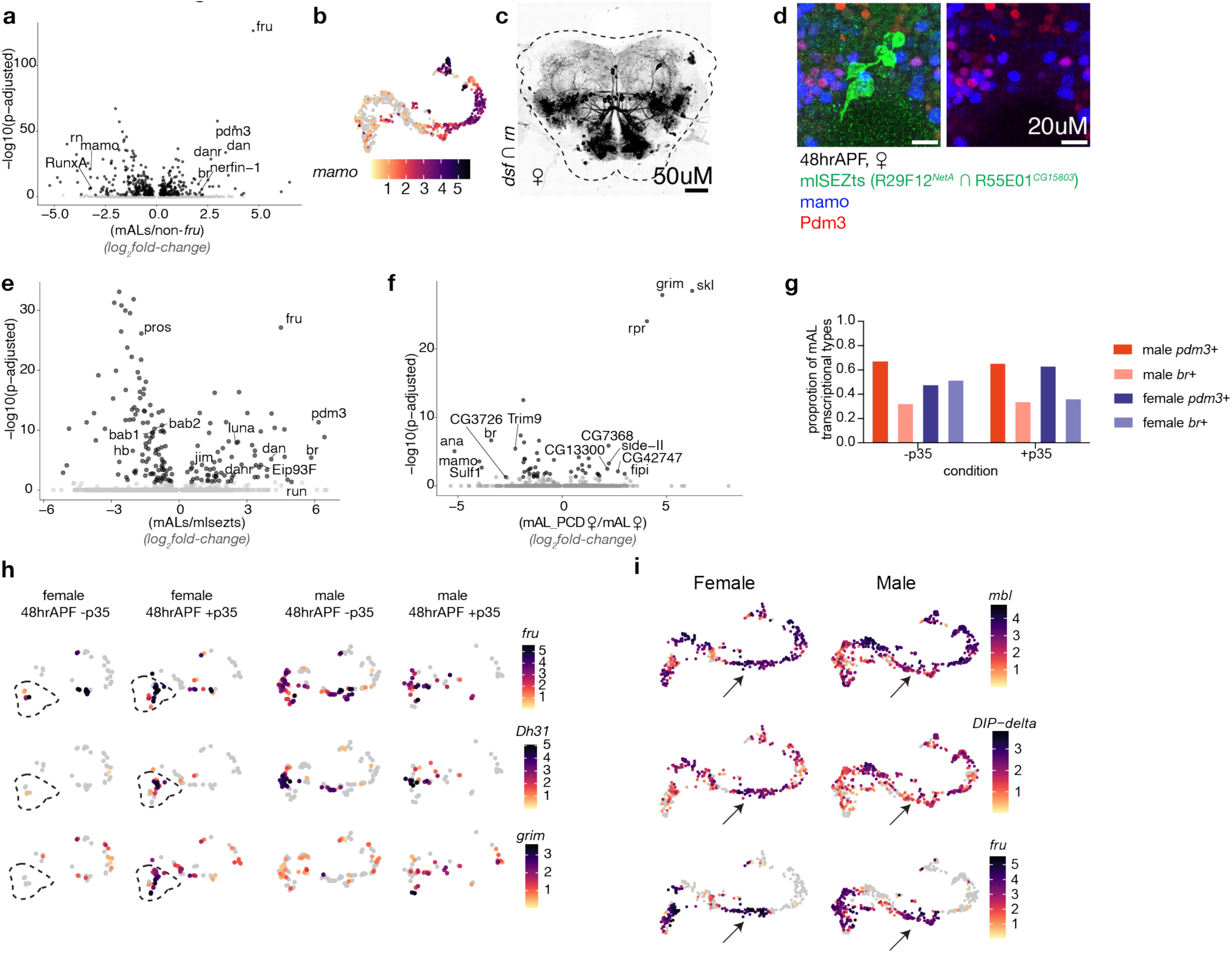
Additional analyses of gene expression differences among CREa1B subtypes and across sexes. **a**, Volcano plot showing differential gene expression between *fru* mAL’s and non-*fru* CREa1B subtypes. **b**, UMAP plot showing log-normalized expression of *mamo* in subclustered CREa1B neurons. **c**, *rn*-expressing CREa1B neurons of females have complex dendritic wings; male sample shown in Figure 6. **d**, Histology for Pdm3 and Mamo protein on the mlSEZt splitGAL4 genotype. Most of the cells labeled in this genotype have small soma and the y-shaped dendrite and ring-shaped axon. The small soma lack both Pdm3 and Mamo protein. The single large somata per hemisphere expresses Mamo. As neurons with larger soma are usually early-born or primary neurons, this somata most likely corresponds to a single winged CREa1B neuron per hemisphere that is also included in the mlSEZt splitGAL4 expression pattern. **e**, Volcano plot showing differential gene expression between mAL and mlSEZt neurons. **f**, Volcano plot showing differential gene expression between “natural” female mAL neurons (i.e. mAL*^br^*) and resurrected female cells that usually undergo programmed cell death (i.e. mAL*^pdm3^*). **g,** Proportion of the *br* and *pdm3* mAL subtypes in male and female, with and without p35. **h**, Dh31 expression is male-specific because it is restricted to the pdm3 temporal cohort, which apoptoses in females. **i**, Examples of genes that are differentially expressed in male versus female mAL neurons.

